# Characterization of non-human primate dura in health and neurodegeneration

**DOI:** 10.1101/2022.06.03.477517

**Authors:** Christopher Janson, Kyle Hauser, Scott Muller, Eric Hansen, Cornelius Lam, Jeffrey Kordower, Liudmila Romanova

## Abstract

Brain meninges and associated vasculature participate in brain clearance and are implicated in many neurological conditions such as Alzheimer’s disease, meningitis, multiple sclerosis, stroke, and traumatic brain injury. However, most of our knowledge concerning brain clearance via meninges is based on rodent data, and relevance to human disease remains unclear. One of the technical barriers in studies of meningeal physiology in health and disease is that existing imaging modalities are suboptimal for large and optically non-transparent meningeal tissue of humans and non-human primate (NHP) animal models. To address this barrier, we performed first characterization of NHP dura by high resolution confocal microscopy of clarified tissue. We investigated vascular structures and resident cells in normal monkeys and primate models of tauopathy and synucleinopathy. We demonstrated the presence of an extensive meningeal vascular network covering the entire tissue surface with resolution to the smallest capillaries. This is also the first work to map lymphatic vessels in the dura of non-human primate (NHP). Overall, the NHP lymphatic meningeal system resembles the anatomy found in rat dura, but it is more complex. Analysis of dura from NPH models of tauopathy and synucleinopathy revealed an association with disease-specific biomarkers (amyloid, tau, α-synuclein) with both the blood and lymphatic vasculature. This work has broad relevance to many brain diseases where solute accumulation and abnormal macromolecular clearance is a part of the pathogenesis.

## Introduction

Significant progress has been made in recent years in our understanding of the diverse roles of brain meninges, including their contribution to neurological disorders.^1–9^ The most obvious and well-known function of the meninges is support and protection of the central nervous system (CNS), achieved by enveloping it and creating space for cerebrospinal fluid (CSF) to buffer and insulate underlying structures. Meninges create a tight chemical-physical barrier between brain fluids and the peripheral circulation, known as the meningeal CSF-blood barrier.^1,4^ This anatomical position and unique structural features of meningeal tissues prevent free exchange of macromolecules and cells between CNS and other fluids and organs. In addition to a barrier function, meninges play a role in brain development and brain cellular homeostasis, harboring stem cells and regulating immune cell trafficking into CNS.^10–12^ These functions make meninges important for brain diseases involving a compromised barrier or immunological components such as Alzheimer’s disease, meningitis, multiple sclerosis, stroke, and traumatic brain injury (TBI).^10,11,13–17^ Understanding meningeal barrier structure, particularly the role of associated vasculature and resident immune cells, is relevant to many neurological conditions.

Meninges (dura, arachnoid, pia) are layered membranes with a distinctive cell and matrix composition along with an integrated blood and lymphatic system. Dura is juxtaposed to the cranium and is comprised of extracellular matrix containing mainly collagen.^18^ In primates, the dura is relatively thick, over 500 µM, and has a skin-like appearance.^19^ Dura in humans and other primates is highly vascularized and innervated, and harbor a diverse population of resident immune cells.^7,20^ Large veins known as dural venous sinuses are folded within the dura and mediate venous outflow from the brain.^4,19,21^ In humans and other primates, the dural layer can be identified visually and may be dissected from inner layers of arachnoid and pia. Such separation is not possible in rodents, in which meningeal layers (particularly arachnoid and dura) are much more fragile and appear as one transparent membrane. Arachnoid and pia together comprise the leptomeninges. Arachnoid interfaces CSF in the subarachnoid space (SAS) overlying the brain surface, and forms trabeculae with blood vessels and CSF cisterns around the brainstem. Loosely adherent to the inner aspect of dura, the arachnoid forms a barrier between CNS and fenestrated blood vessels of the dura, which are accessible to peripheral circulation. Epithelial cells of arachnoid are connected with tight junctions and desmosomes which comprise the CSF-blood barrier, regulating transport between the SAS and dural veins. ^22–24^ Arachnoid membranes themselves are largely avascular, though they surround traversing vessels in the SAS. In contrast with arachnoid, pia is relatively permeable to molecules as large as albumin.^25^ Pia interfaces the glia limitans formed by astrocyte end feet and invests vessels as they penetrate the brain.

Meninges have a complex morphology with many involutions. Major meningeal folds that we will consider in this manuscript, which harbor a variety of critical draining vessels, are falx cerebri and tentorium cerebelli of the dura. In the midline, the dura invaginate to follow the longitudinal fissure between the cerebral hemispheres, and this dividing structure is known as falx cerebri. In primates, the falx cerebri accommodates a large superior sagittal venous sinus. Tentorium cerebelli is another major fold, which forms the roof of the posterior cranial fossa separating the occipital and temporal cerebral hemispheres from the cerebellum and infratentorial brainstem. Both falx cerebri and tentorium cerebelli maintain the anatomy of the brain by supporting its weight and providing a matrix for attachment of blood vessels.^26^

Currently, our knowledge about meninges is rapidly changing especially in terms of their role in brain immunity and solute efflux from the CNS.^4,7,27^ Given their relatively superficial location and rich vascular structure, meninges are a potential therapeutic target for neurodegenerative disorders in which metabolic waste is removed from the brain. In developing strategies to regulate transport properties, it is important to understand meningeal structure, especially in humans and non-human primate (NHP) models. However, our knowledge comes primarily from readily available rodent models. There is high variety of surgical and imaging approaches for rodents and rodent meninges are more accessible to the studies due to their smaller size. Meninges in humans and larger animals present challenges because of the limited experimental toolbox and access. Moreover, larger tissue area of the meninges in primates significantly limits imaging modalities. Since meninges are flat and multi-layered structures, spatial relationships among different components can only be adequately assessed in whole-mount preparations. This approach has been used in rodent models with smaller and optically transparent meninges.^28–30^ However, imaging remains challenging in primates due to the tissue volume and opacity. Investigation of NHP meninges in earlier studies was based on injections of different substances such as carbon particles or dyes into the subarachnoid space, followed by analysis with light or electron microscopy.^31,32^ This provided either an overly cursory view of the tissue in the case of LM, or an overly limited view with TEM. With human tissue, whole meninges are difficult to obtain, which results in a circumscribed anatomical context. Most human preparations have traditionally been small paraffin embedded fragments, sectioned perpendicular to the tissue plane. This approach provides a limited perspective and omits three-dimensional vascular relationships.^16,33^ It is particularly limiting in showing only the vessel lumen without critical information on other characteristics of vascular structures such as size, branching patterns, and total cross-sectional area. This limitation has become particularly apparent in studies of meningeal lymphatic vessels when substantial efforts were devoted to establishing their existence in primates. Several groups reported detection of meningeal lymphatic vessels using non-invasive human imaging but those modalities do not provide sufficient resolution for precise identification of vascular relationships.^34,35^ Prox-1 and podoplanin (PDPN) positive vessels were identified in marmoset monkey dura in the areas of superior sagittal sinus (SSS) and middle meningeal artery (MMA), but details of the vascular network are not described.^35,36^

Here we report the first characterization of non-human primate dura using high resolution confocal microscopy of clarified tissue. We characterize spatial relationships among vascular structures, resident immune cells, and extracellular matrix in normal monkeys and vasculature in models of tauopathy and synucleinopathy. We demonstrate the presence of an extensive vascular network covering the entire tissue with resolution to the smallest capillaries. This is also the first work to map meningeal lymphatic vessels alongside extensive venous networks in the dura of non-human primate (NHP). We found that lymphatic vessels can be separated into two types based on localization and morphology. The first type clusters along major venous sinuses and appear as blind-ended vessels. The second type is localized laterally alongside the MMA and venous branches following the arteries. Overall, the primate lymphatic system resembles rodent system, yet it is not identical. In our analysis of dura from NPH models of tauopathy and synucleinopathy, we found disease specific biomarkers (amyloid, tau, α-synuclein) associated with both blood and lymphatic vasculature, suggesting that both may be involved in metabolic clearance. Our protocol for acquisition of high quality images in intact dural tissue also provides spatial information on the population of resident cells such as macrophages and fibroblasts as well as extracellular matrix. The imaging modalities described here will be useful for further analysis of dura in NHP and human tissues.

## Results

In this manuscript we report characteristics of the dura mater of Cynomolgus monkeys (Macaca fascicularis) and key anatomical and pathological features of dura in models of tauopathy in Rhesus monkey (Macaca mulatta) and synucleinopathy in Cynomolgus monkey.^37,38^ Using whole-mount preparations of the entire tissue in combination with high-resolution confocal microscopy, we investigated morphology and localization of vascular and lymphatic beds, resident cells and extracellular matrixes. For models of proteinopathies, we also investigated distribution of the deposition of clinically relevant proteins such as amyloid (Aβ), tau, and α-synuclein.

### Monkey dura is highly vascularized

Dura was removed during necropsies and in most cases cut only in half along the midline or in some cases the area of falx cerebri was dissected separately from the lateral areas. For larger specimens, we sometimes cut into 4 sections with the halves cut into frontal and dorsal sections. Tentorium cerebelli was dissected as a separate piece. Unlike rodent dura, monkey dura resembles human tissue and is highly opaque (Figure 1, A). Therefore, it was critical to develop clearing protocol and all our specimen preparation included a clearing step that rendered the tissues optically transparent (Figure 1, B and C). The first step for us was to assess the structure of the vasculature in its entirety in the whole-mount preparation of the dura. For visualization of blood vessels, we used CD31 as a primary marker for blood endothelial cells.^39^ Staining with CD31 antibody revealed a dense and extensive network of arteries and veins, as well as capillaries and venules covering the entire dura as shown on the scans of lateral dura and tentorium (Figure 2, A and B). Arteries, including large branches of the middle meningeal artery (MMA) were clearly identifiable as marked on Figure 2, A. For simplicity, we will refer to MMA and all of its divisions as MMA. Large and smaller meningeal arteries were always flanked by veins (Figure 2, C and D). We also frequently observed groups of round CD31 positive cells freely distributed without association with blood vessels. Falx cerebri has a particularly complex venous drainage network as seen in (overview) Figure 3A and (close up view) Figure 3B. In most cases, it was possible to clearly distinguish veins and arteries based on location and morphology. To verify this observation, we performed staining with smooth muscle actin (SMA) that distinguishes arteries from veins.^40^ The majority of falcine blood vessels were confirmed as veins (Figure 4B, C). Typically, large and medium size veins flank large arteries and smaller arteries may run parallel with only one vein (Figure 4B).

**Figure 1.**
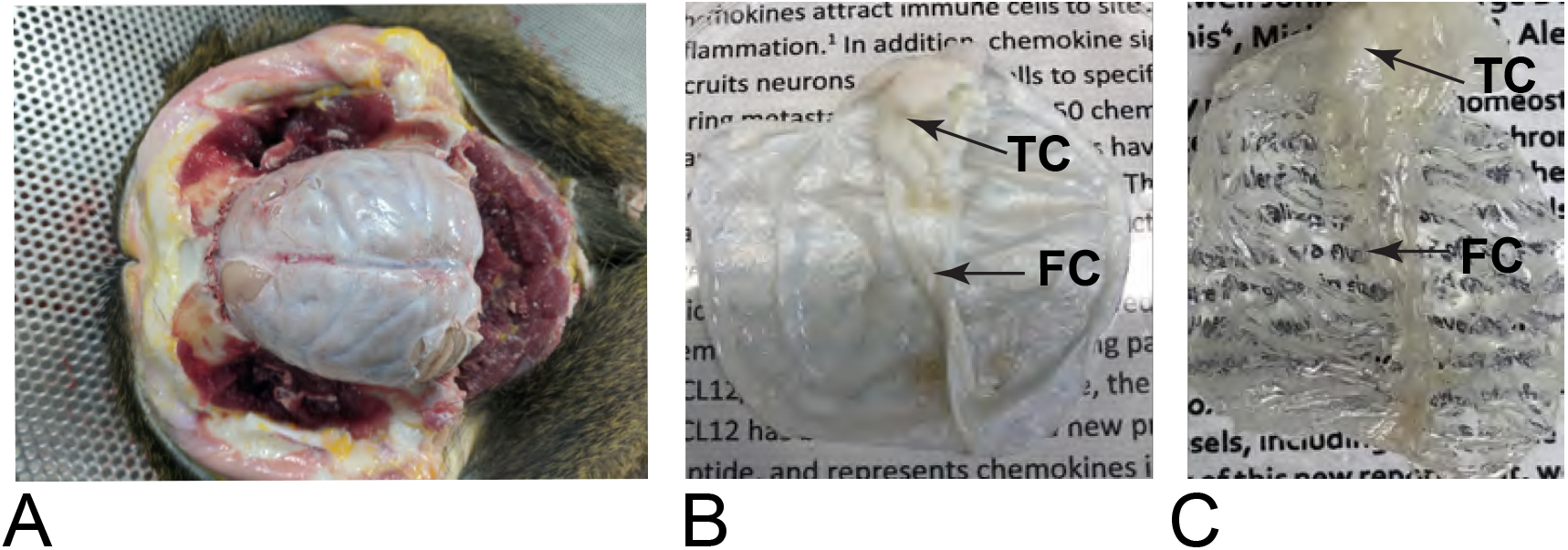
Cynomolgus monkey dura is large, thick and optically non-transparent tissue. Cynomolgus monkey dura during dissection (A), before clearing (B) and after clearing (C). FC – falx cerebri, TC – tentorium cerebelli.

**Figure 2.**
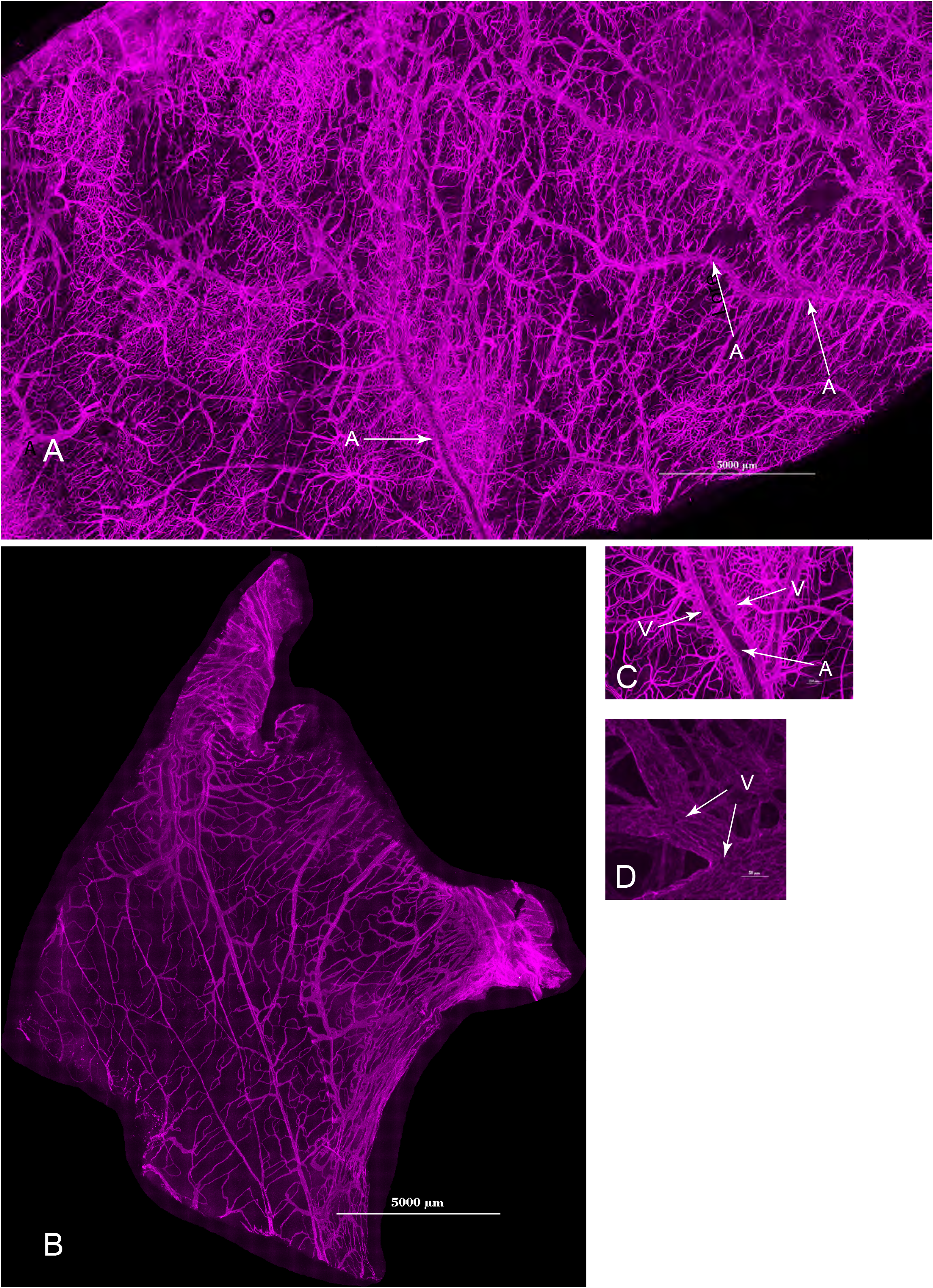
Dura is highly vascularized in lateral areas and tentorium cerebelli. Panoramic view of monkey dura probed with CD31 antibody to visualize blood vessels. (A) Lateral area of the dura cut along falx cerebri and a portion of tentorium cerebelli (B). A panel of the representative close up fields of the vasculature in the lateral area showing arteries, veins and capillaries (C, D). V – veins, A – arteries, C – capillaries. Scale bars are 5000 µm on (A) and (B); 250 µm on C, 50 µm on D.

**Figure 3.**
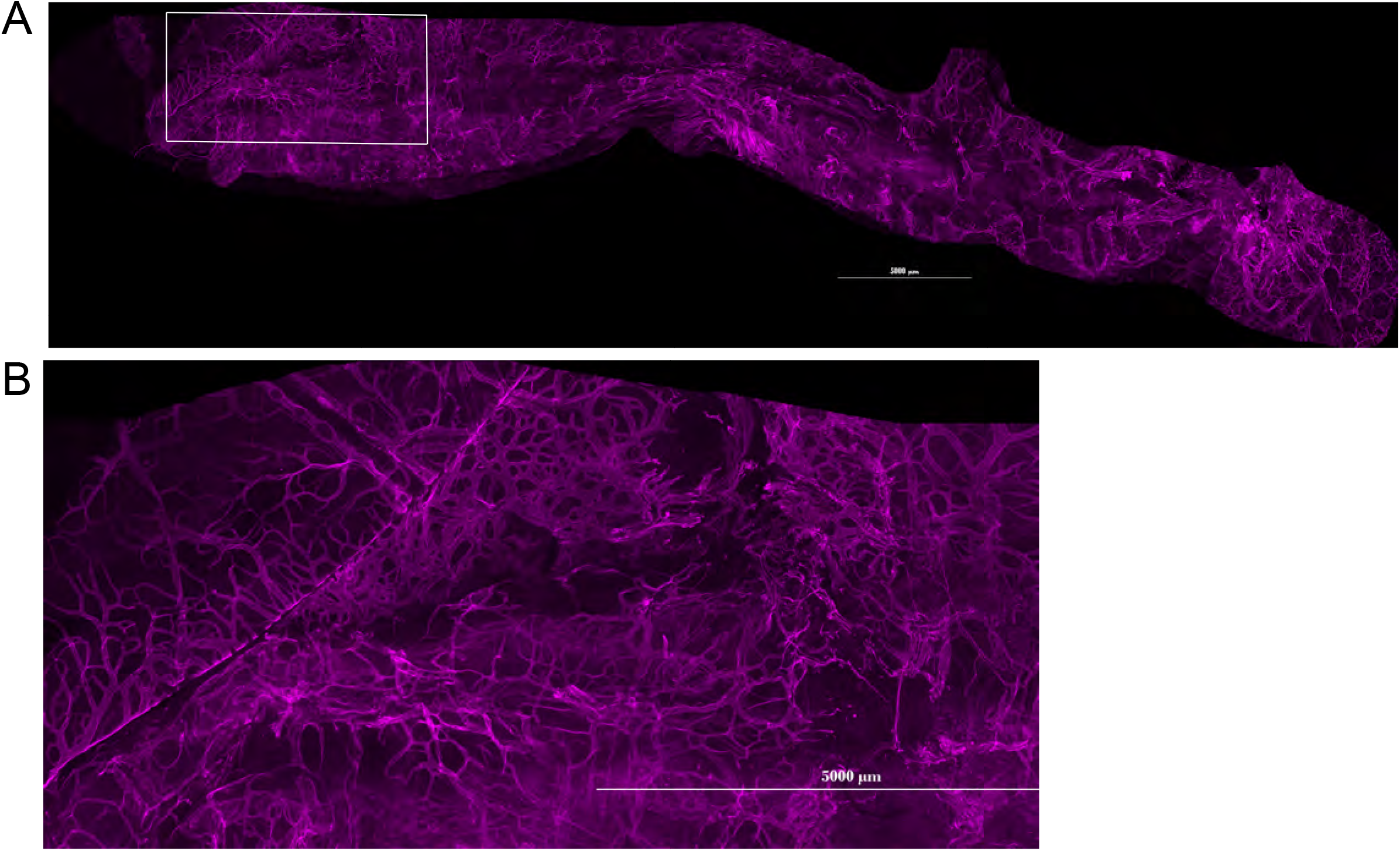
Blood vessels of the falx cerebri. (A) Overview of the dissected area of the flax cerebri with dense network of blood vessels shown. (B) shows close up view of the area in rectangle on (A). Scale bars are 5000 µm on (A) and (B).

**Figure 4.**
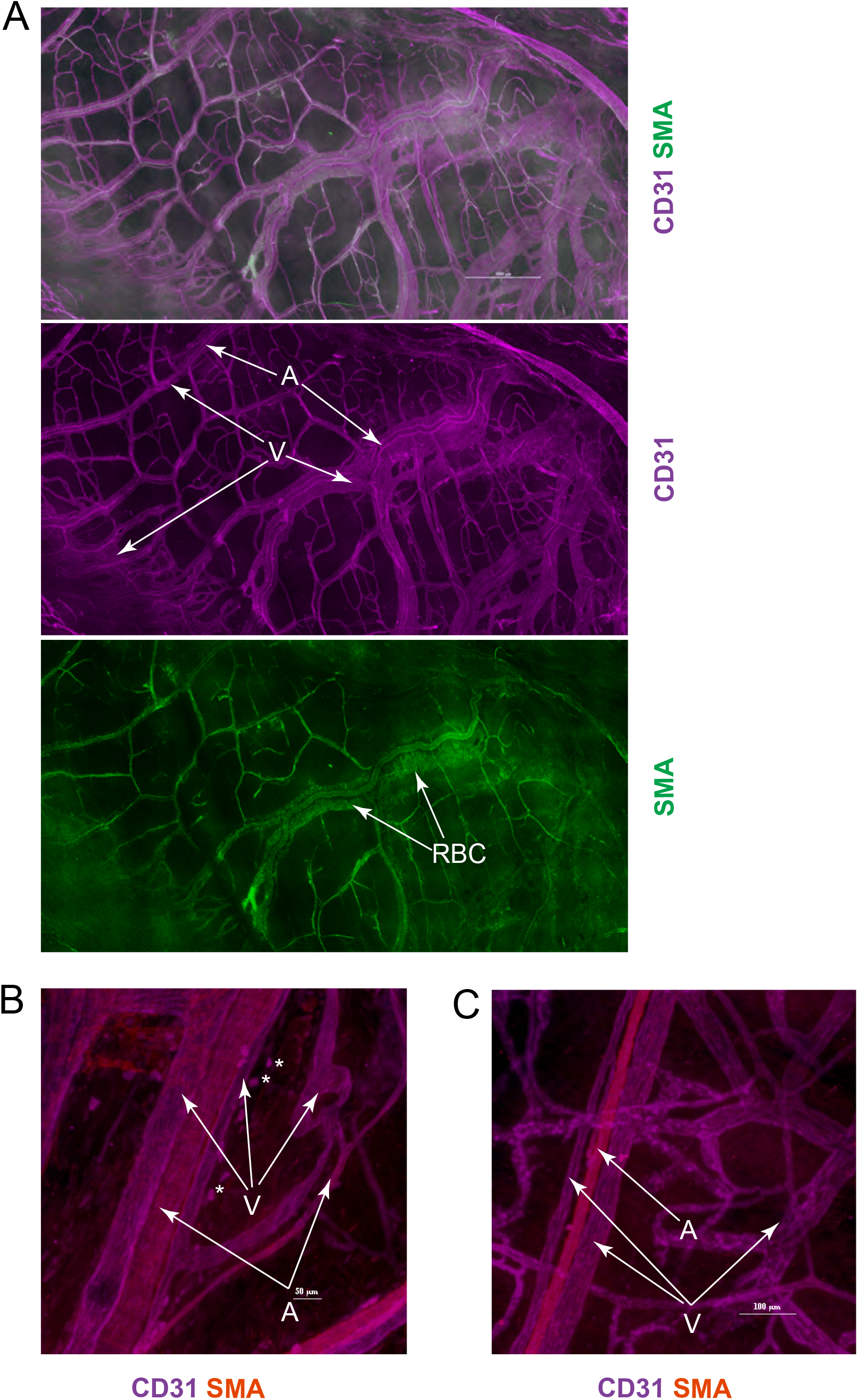
Venular structures dominate vascular network in monkey meninges. Staining with SMA allowed to visualize arteries in meningeal vascular network. Substantially larger veins contain residual blood seen on green channel (A, arrows) and showing typical position of arteries flanked by veins. Representative images of anatomy are shown on B-C. Scale bars are 1000 µm on (A), 50 µm on (B) and 100 µm on C. V – vein, A -artery. Asterisks indicate CD31+ cells associated with blood vessels.

### Lymphatic vessels in monkey dura are clustered around falx cerebri and run laterally along MMA and branching arteries

The presence of lymphatic vessels has been demonstrated in dura in mice and rats.^28–30,36^ There is limited evidence that a similar system exists in primates and humans.^16,33,35,36^ Staining with antibodies against LYVE-1 (hyaluronan receptor-1), PDPN (D2-40/podoplanin), and Prox-1 (Prospero Homeobox 1) is conventionally used for identification of lymphatic endothelial cells.^41,42^ Other indicators of lymphatic phenotype include expression of 5′-nucleotidase (5′-Nase), which is also highly expressed in arachnoid, and reactivity to LA102 antibody.^43,43–45^ We found strongly PDPN positive vascular structures located in two areas of dura. First, in a branched network in the falx (Figure 5, A). Second, in paired vessels running laterally along veins flanking MMA and its branches (Figure 15 and 16). These two types of PDPN+ vessels have different morphology and density. We will refer to them hereafter as lymphatics based on their position and expression of PDPN marker. The density of lymphatics is high around SSS and the network consists of blind ended vessels that extend laterally from the falx (Figure 6, A). We frequently observed areas where the lumen was enlarged forming structures that appear as “bulbs” (Figure 6, B). The diameter of the “bulbs” varied between 100 to 300 µM. It is unclear if this is partially an artefact of tissue processing, but these “bulbs” were only observed in SSS lymphatics. PDPN+ vessels that run along MMA do not have extensive branching. The branching in these vessels is limited to the area of arterial anastomoses where lymphatics continue to follow the arteries (Figure 16). Intensity of PDPN staining was consistently lower in these vessels than in the vessels around SSS.

**Figure 5.**
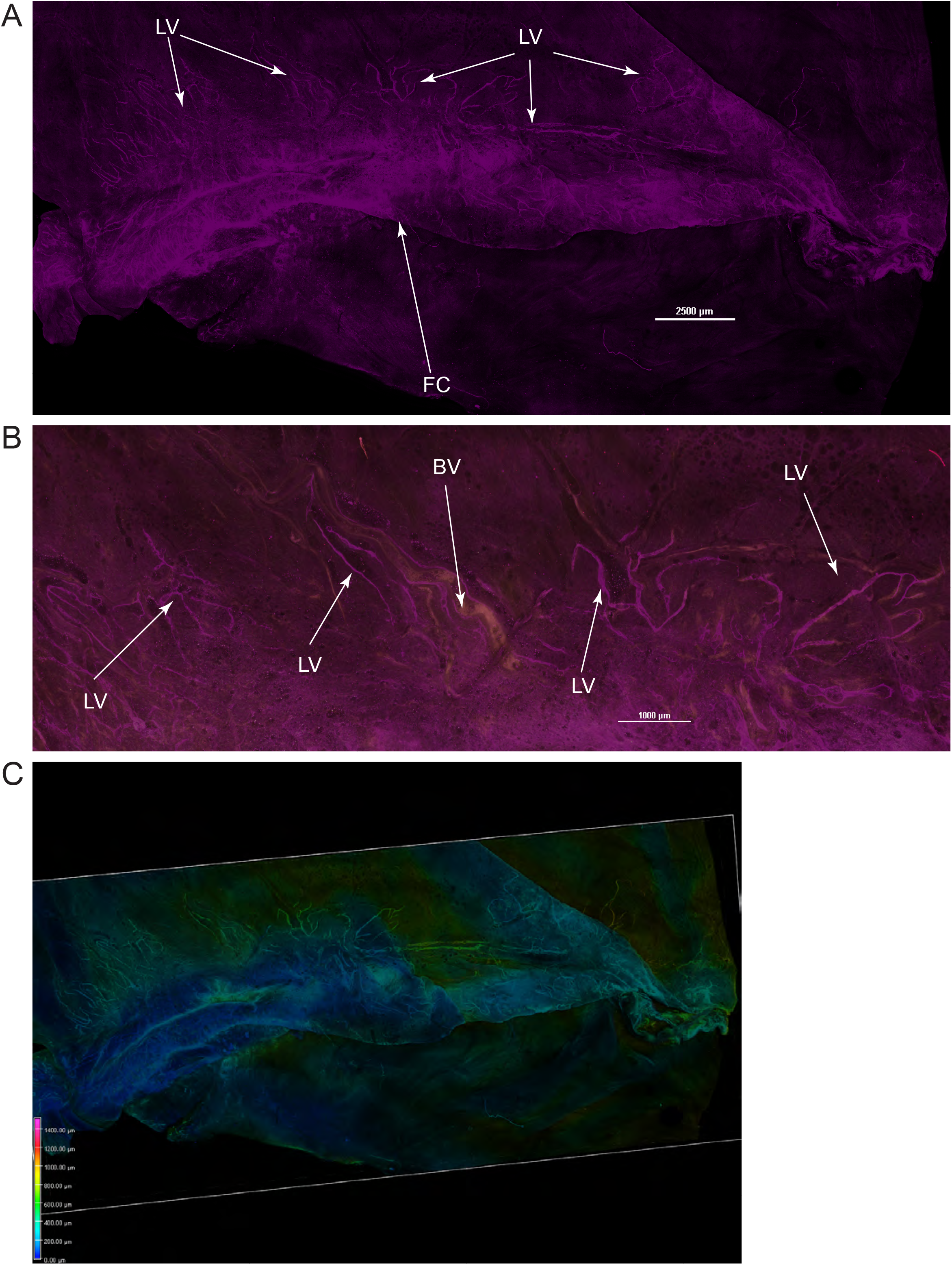
Lymphatic vessels of the falx cerebri. (A) PDPN+ vessels identified in the area of falx cerebri in the proximity to SSS as seen on whole-mount monkey dura. (B) Close up view of the region shown on (A). (C) 3D depth projection of the image on (A). LV – lymphatic vessel, BV – blood vessel. Scale bars are 2500 µm on (A) and (C) and 1000 µm on B.

**Figure 6.**
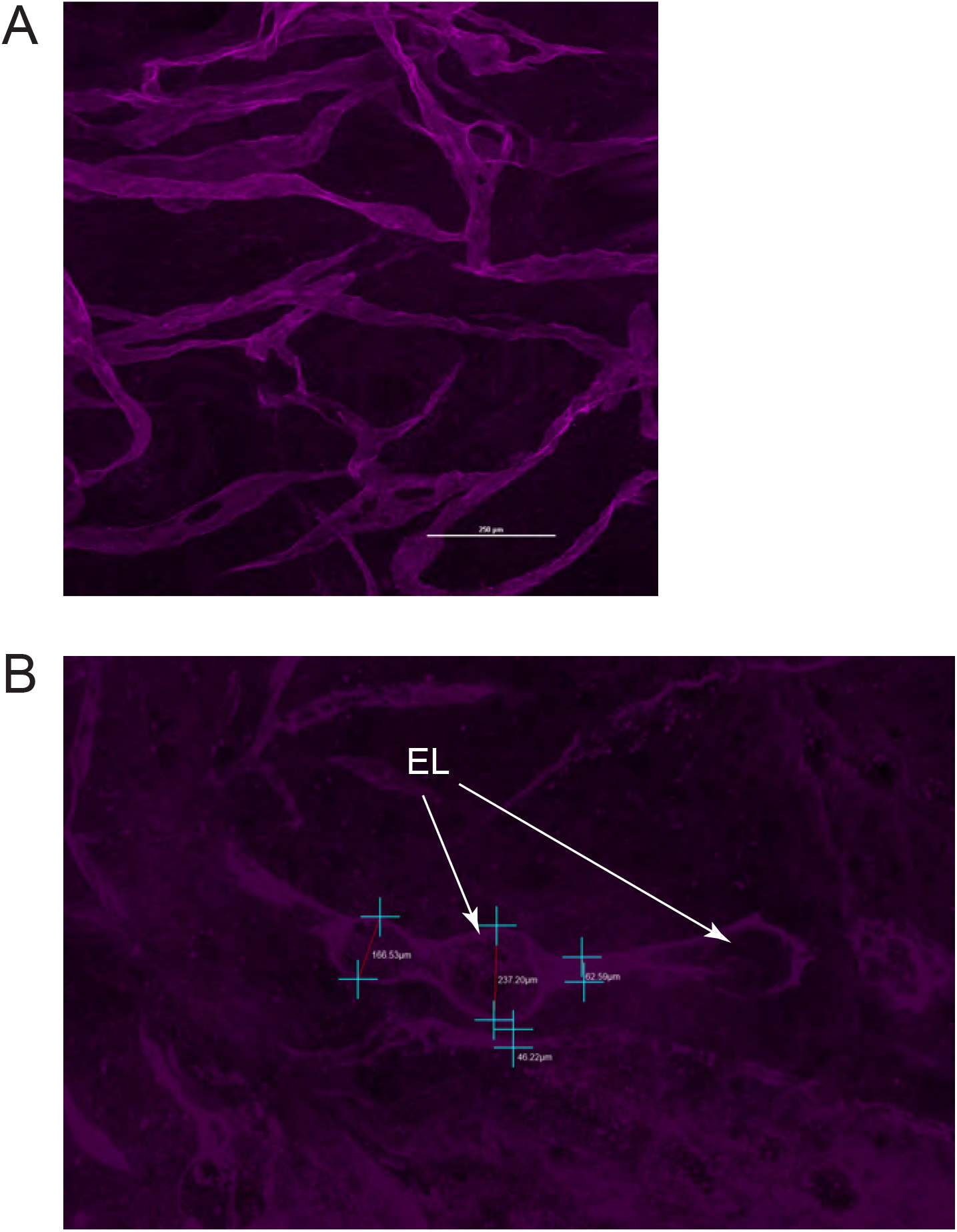
Close up view of the SSS lymphatics. (A) PDPN+ lymphatics extending from the area of falx cerebri appear as blind ended vessels. (B) Enlarged lumen areas in the SSS lymphatics. EL – enlarged lumen. Scale bar is 250 µm on (A). Measurement markings on (B) demonstrate potential variations in the lumen diameter.

Staining with LYVE-1 antibodies showed that PDPN+ vessels were only weakly positive for LYVE-1. However, we observed LYVE-1+ structures densely covering almost the entire dura with high density around large veins (Figure 7). In high magnification imaging with 20x and 40x long working distance lenses, these LYVE-1+ structures appear as a network of elongated processes resembling lymphatic capillaries (Figure 8, A) and had very weak PDPN staining (Figure 8, B). It was difficult to assess connections between individual vessel-like structures with Maximum Intensity Projections of very thick tissues. Volume projection of z-stacks showed that elongated structures are most likely not connected to each other (Figure 8, C). To verify, we attempted tracing of these structures using Vesselucida360 (MBF BioSciences). This software allows reconstruction and modeling of vascular networks. We were not able to reconstruct a continuous network of vessels and follow individual branches within Vesselucida360. While it is possible that LYVE-1 staining is punctate and thus interferes with tracing of capillaries, the most likely explanation is that these lymphatic like structures do not form a proper vascular network. These cells phenotypically resemble spindle shape macrophages, and this prompted us to explore expression of macrophage markers by those cells. We confirmed that LYVE-1 positive cells also expressed macrophage markers as CD206 and IBA.

**Figure 7.**
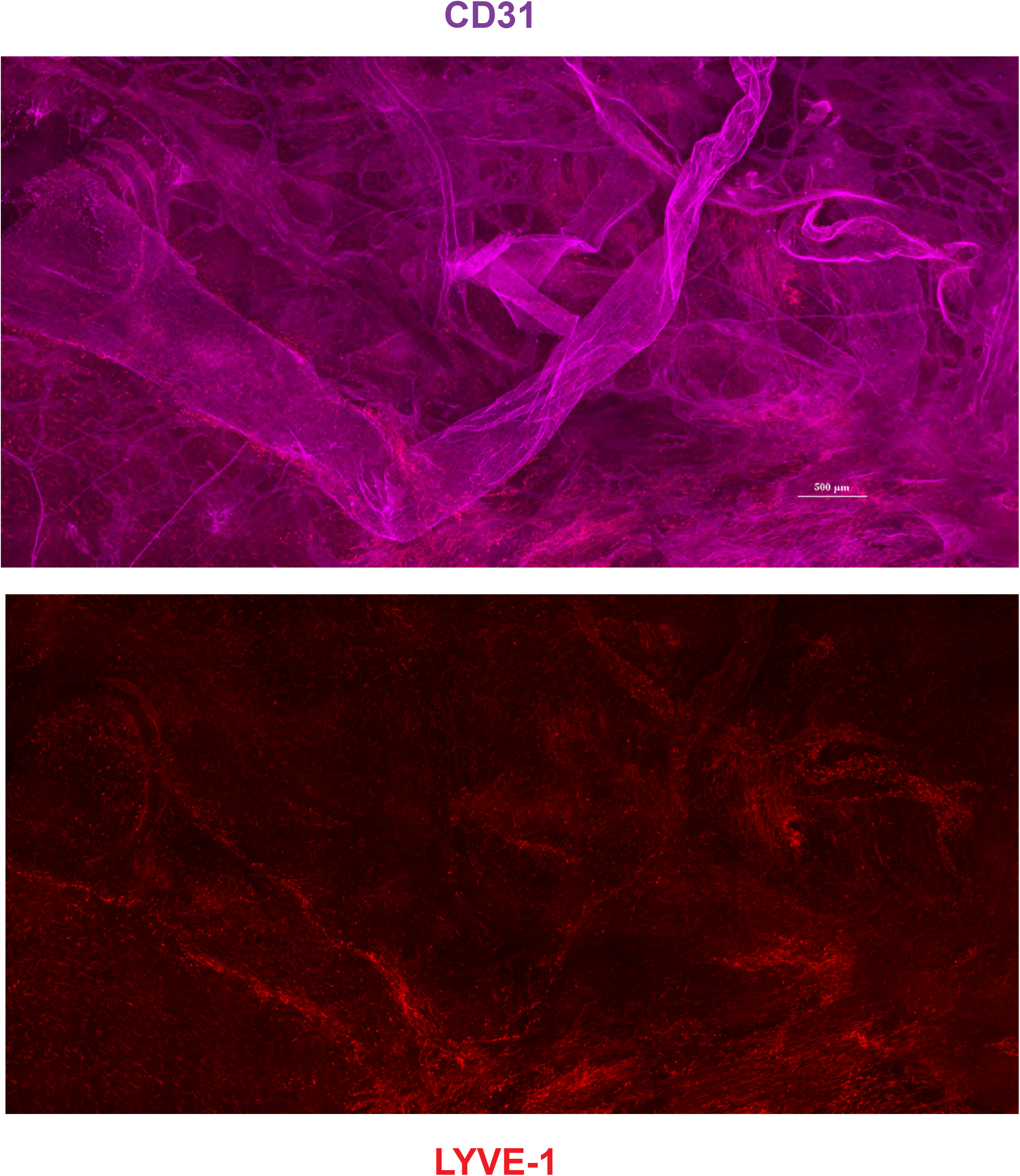
LYVE-1 positive structures in monkey dura. LYVE-1+ structures are seen on this representative image associated with large veins and without association with the vasculature. Scale bar 500 µm.

**Figure 8.**
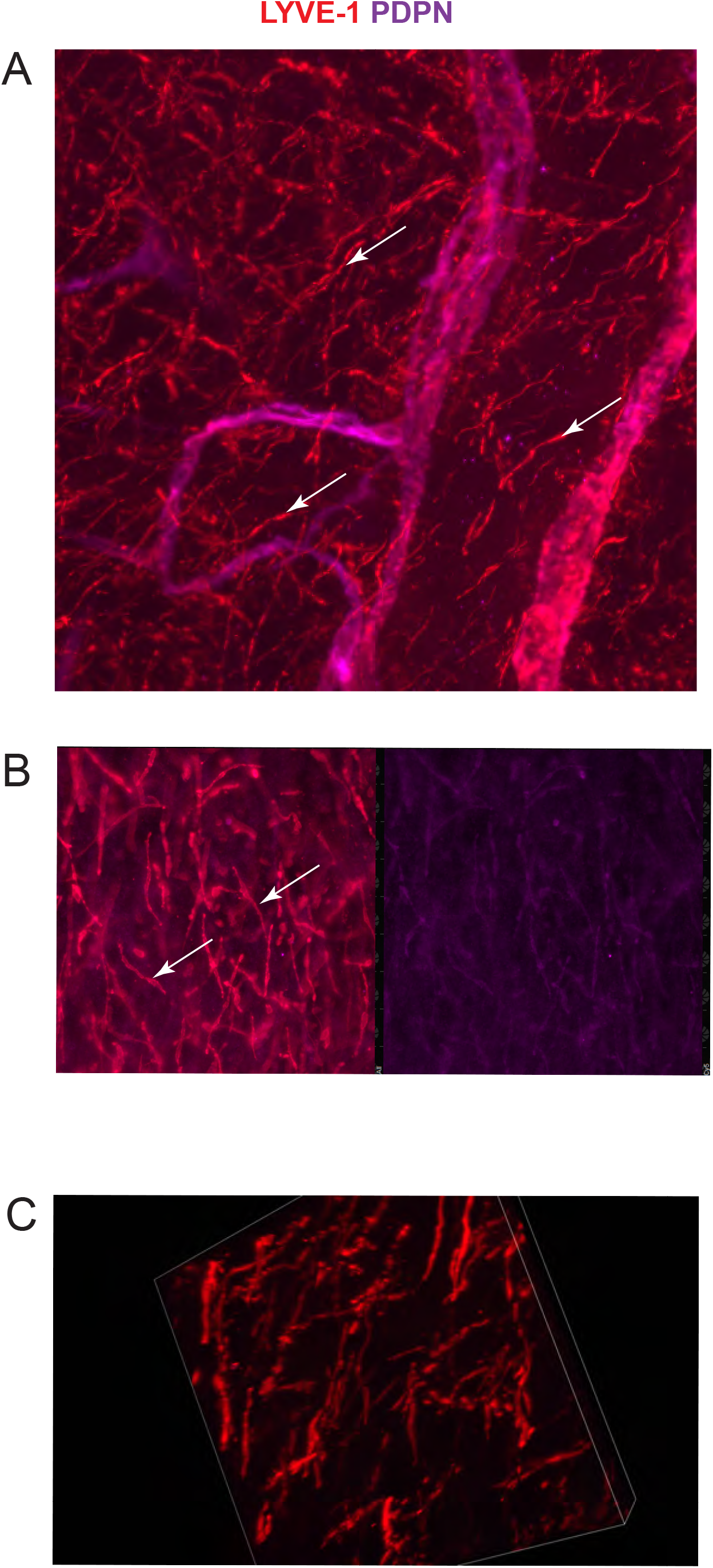
LYVE-1+ macrophages in monkey dura. (A) LYVE+ positive structures of the elongated shape, later identified as macrophages, densely populate monkey dura. (B) LYVE-1+ structures are also weak PDPN positive. (C) 3D projection used for vessel tracing in Vesselucida360. Arrows point on LYVE-1+ structures.

Functionally and morphologically, lymphatic vessels consist of lymphatic capillaries and collecting vessels. Lymphatic capillaries absorb interstitial fluid and collecting vessels propel it towards lymph nodes from where the fluid is returned to venous circulation.^46^ PDPN expression and weaker LYVE-1 expression are indicative of the collecting vessel phenotype. Therefore, we explored that possibility. Collecting vessels are covered with layers of muscle cells, which contain muscle contractile proteins, such as αSMA, and can be used for their identification. Collecting vessels also have valves, which can be identified morphologically with the surface marker staining like PDPN or CD31, or with anti-α-integrin antibodies.^47^ Neither of these approaches were able to identify valves on any of our preparations. Analyzed specimens were also negative for α-integrin. Immunocytochemistry with anti-αSMA antibodies showed that αSMA was only associated with CD31+ blood vessels and not with PDPN+ vessels around the falx, tentorium, and lateral dura (Figure 9 and 10). All PDPN+ and LYVE-1+ structures were negative for 5’-Nase expression and had no reactivity with LA10 antibody.

**Figure 9.**
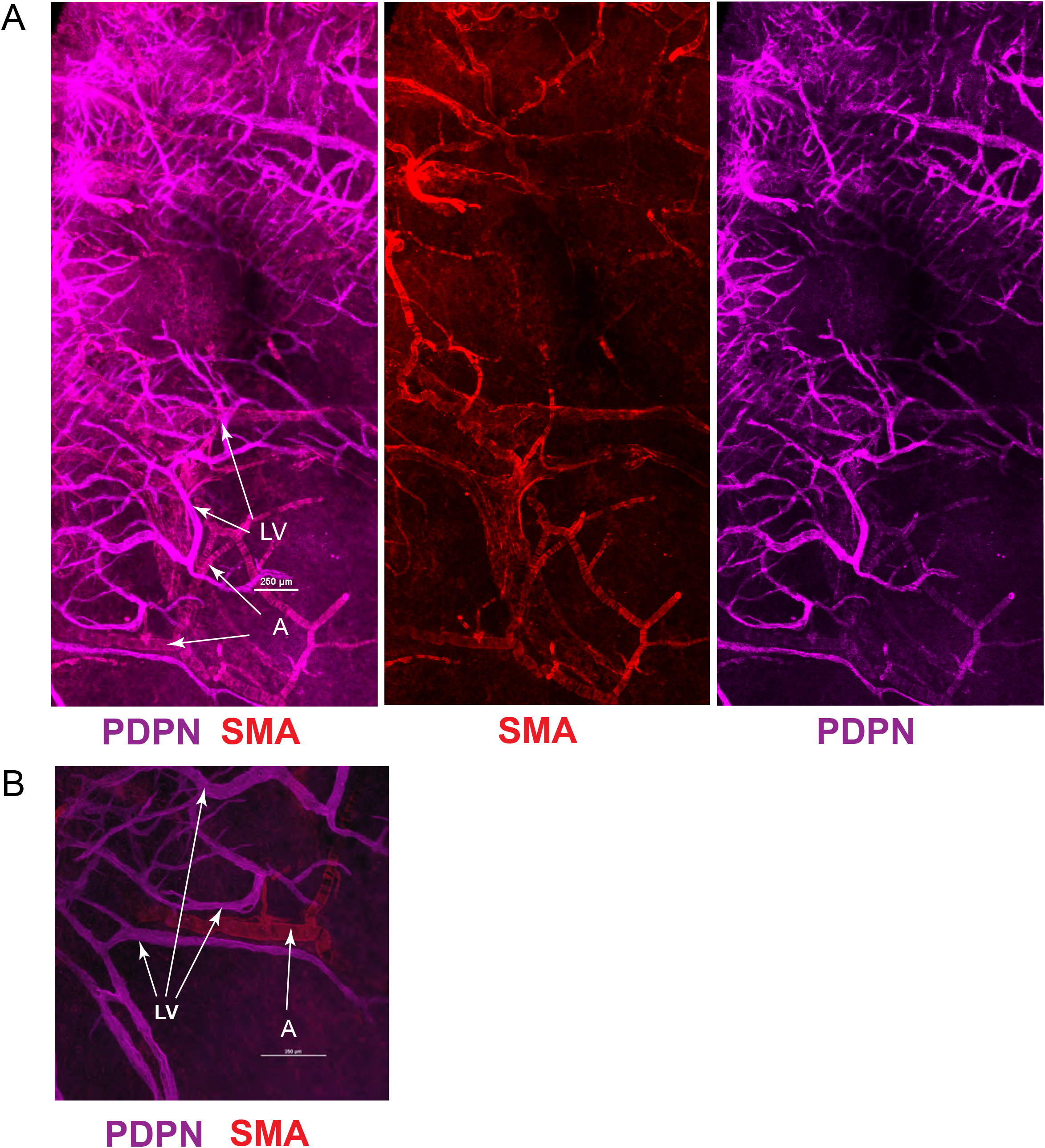
Lymphatic vessels and arteries in the tentorium area don’t have SMA coating. (A) Representative image of PDPN and SMA stained vessels in the area of falx adjunct to the tentorium. PDPN and SMA signals do not overlap and visualize different vessel populations, which correspond to lymphatics and arteries (B). Close up view of frequently observed position of SMA-coated artery in association with lymphatics. LV – lymphatic vessel, A – arterial vessel. Scale bars are 250 µm.

**Figure 10.**
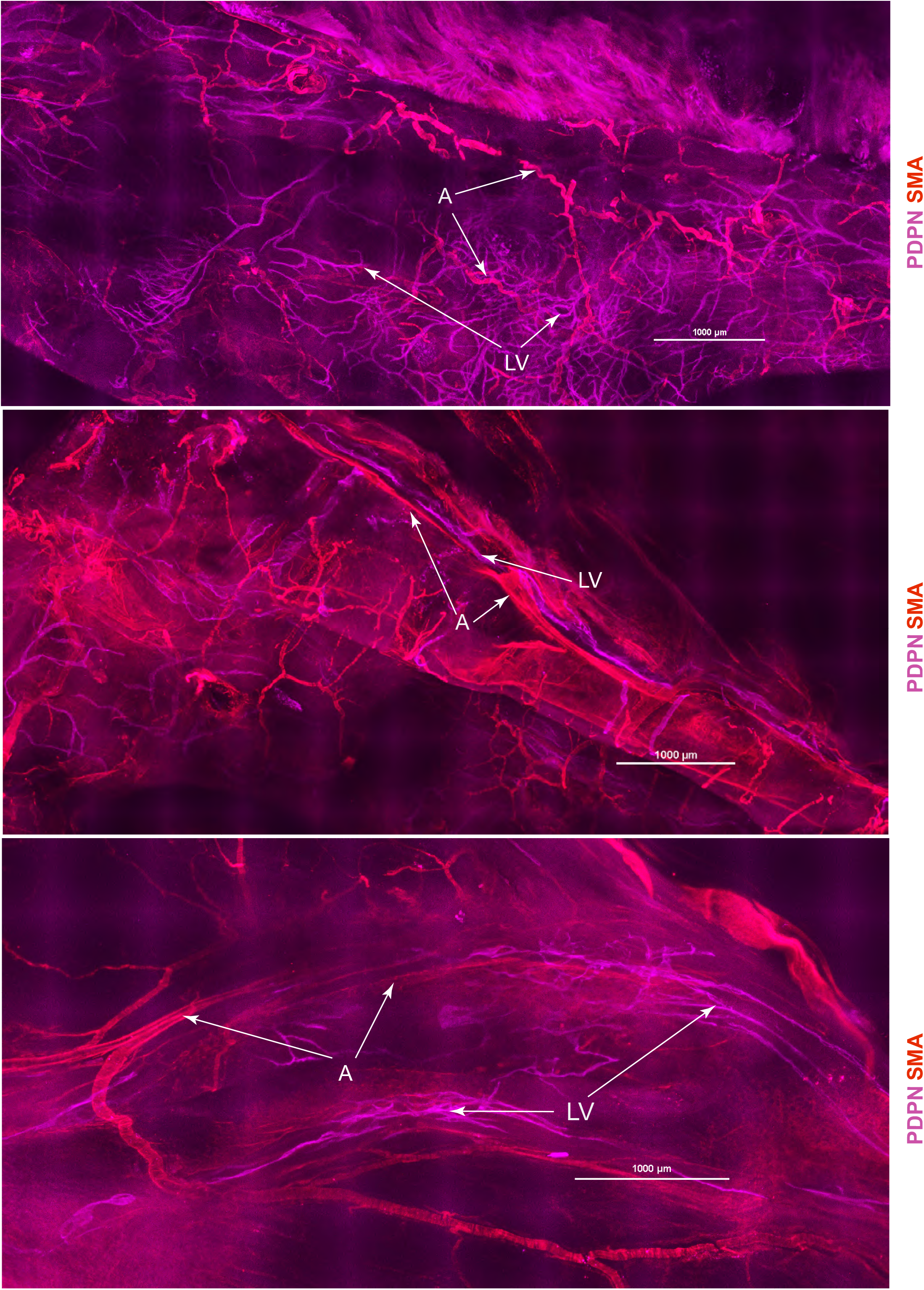
Position of lymphatic vessels and arteries in the falx area. Lymphatics and arteries are visualized with PDPN and SMA staining. LV – lymphatic vessel, A - arterial vessel. Scale bars are 1000 µm.

### Monkey dura is densely populated with CNS-border macrophages

Multiple studies have demonstrated that a variety of cells including macrophages, monocytes, and mast cells reside in the meningeal space.^41^ These cells reside within the meninges, which forms a physical border between the CNS and peripheral circulation. Specifically, macrophages are known to be highly abundant in the dura.^48^ We initially identified a LYVE+ population of meningeal macrophages, and to explore further the presence and distribution of meningeal macrophages, we stained dura with anti-CD206 antibody. CD206 is a mannose receptor which is used routinely as a macrophage marker at the CNS border.^2^ This staining revealed an abundant population of CD206+ cells in the entire tissue as shown in Figure 11 and Figure 12, A. The majority of CD206+ cells have an elongated shape (Figure 12, B and C). There are two different populations; some cells are closely associated with blood vessels (Figure 12, A and C) and another population is individual cells scattered within the entire meningeal surface without association with vasculature (Figure 12, B). We did not identify specific spatial patterns in the position of vessels associated with CD206+ macrophages or in the position of freely residing dural macrophages.

**Figure 11.**
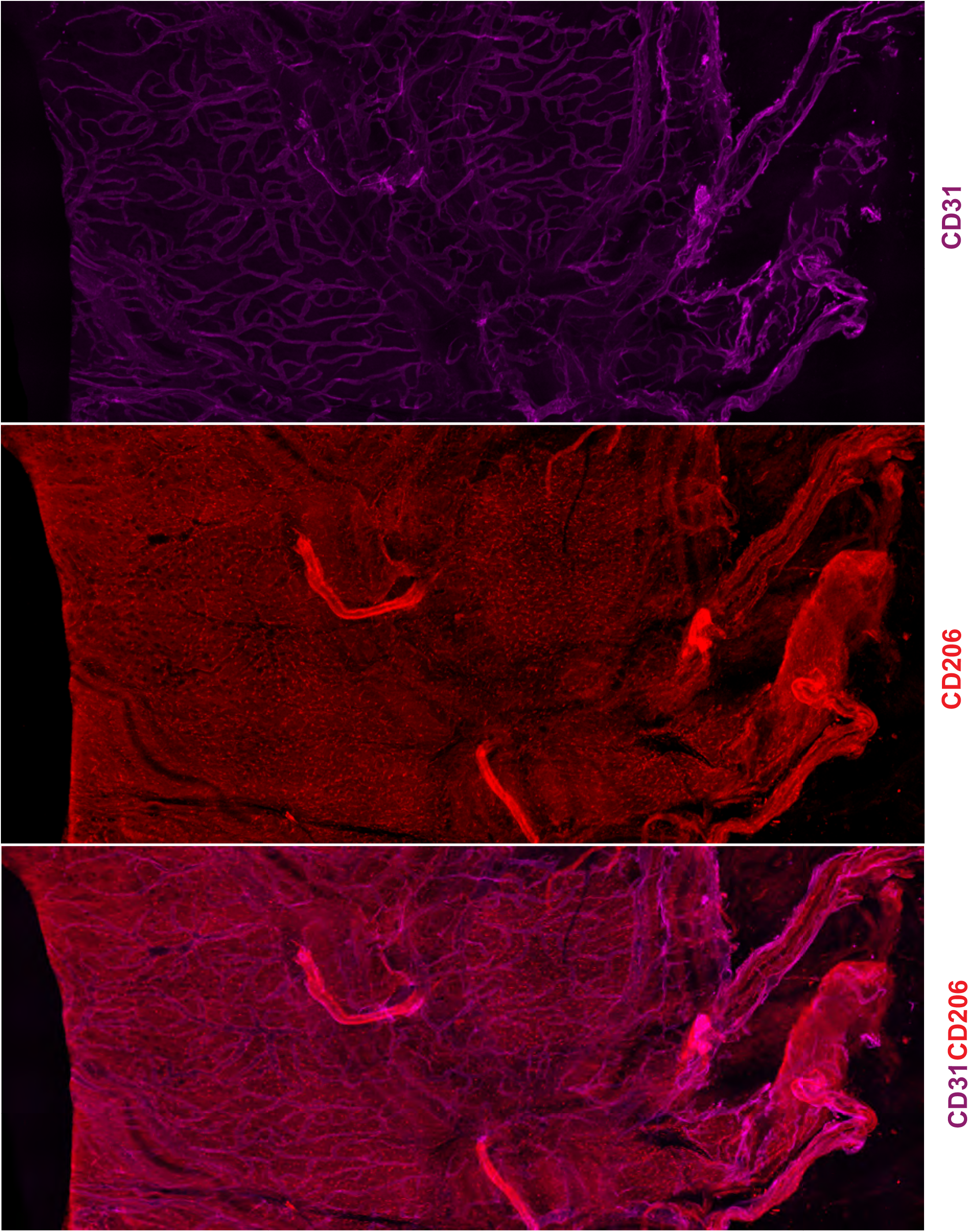
Monkey dura is densely populated with CD206+ macrophages. CD206+ cells align along blood vessels in the adult dura or in the dura without association with vasculature.

**Figure 12.**
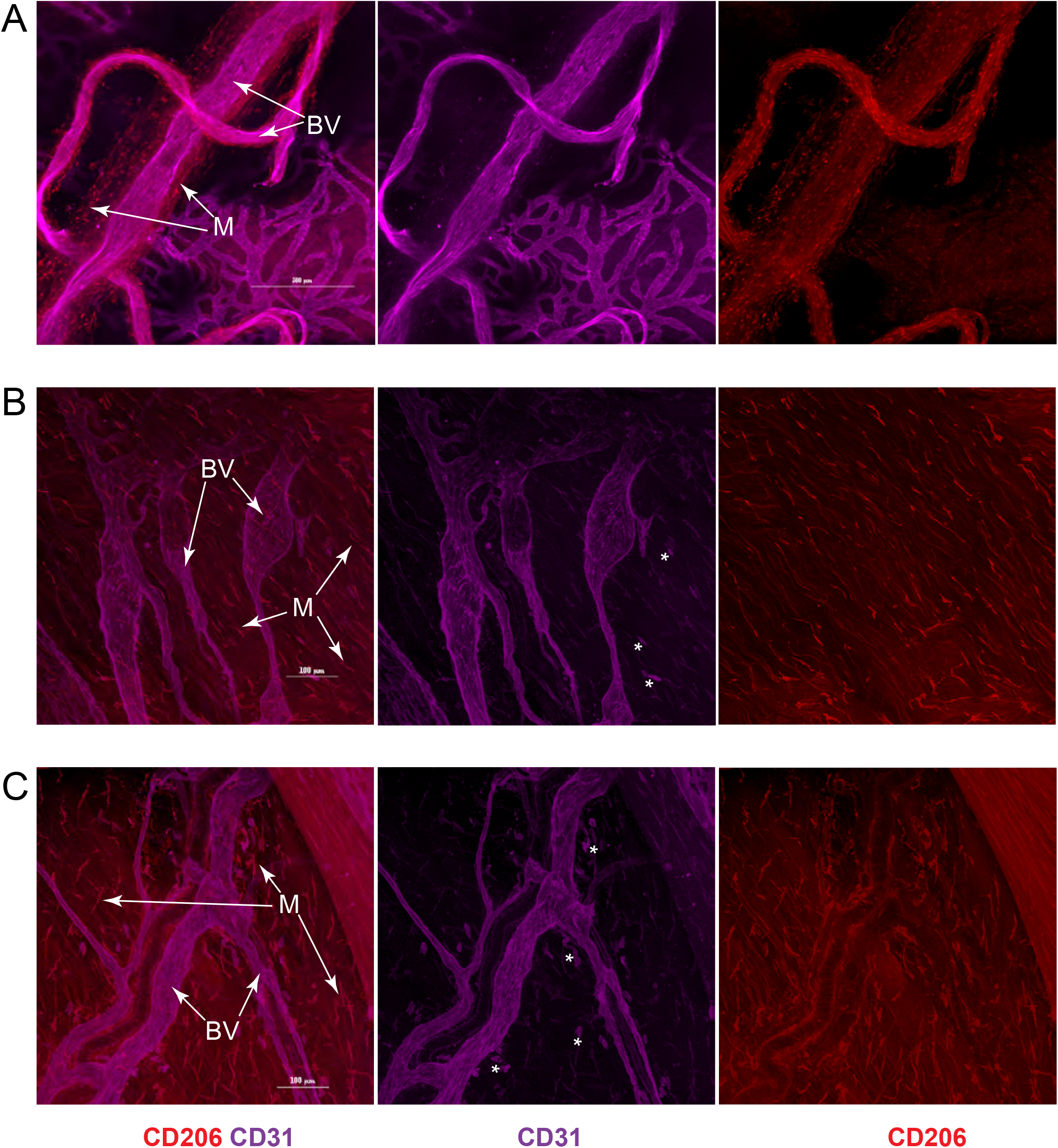
CD206+ macrophage populations are associated with blood vessels or distributed in the dura. (A) shows mostly perivascular localization and (B) primally free cell population. Scale bars are 500 µm on (A) and 100 µm (B) and (C). BV – blood vessels, M – macrophages, CD31+ cells are marked with asterisks.

### Perivascular mast cells reside in the dura

Mast cells are long-term resident cells of the meninges.^6^ Immune resident cells of the meninges such as mast cells appear to play important regulatory roles.^2^ Mast cells reside in border tissues where they involved in rapid immune responses by quickly releasing substances from stored granules including proteoglycans, proteases, leukotrienes, and cytokines.^49^ This is thought to potentiate further immune cell reaction to injury. Probing of dura with CD117 (cKit) antibody showed positive immunoreactivity along blood vessels, primarily of smaller diameter (Figure 13, A). The staining was discontinuous, only in certain areas of the vessels (Figure 13, B). It is known that mast cells also reside in dura without association with the vasculature, but we did not observe that population in our specimens.

**Figure 13.**
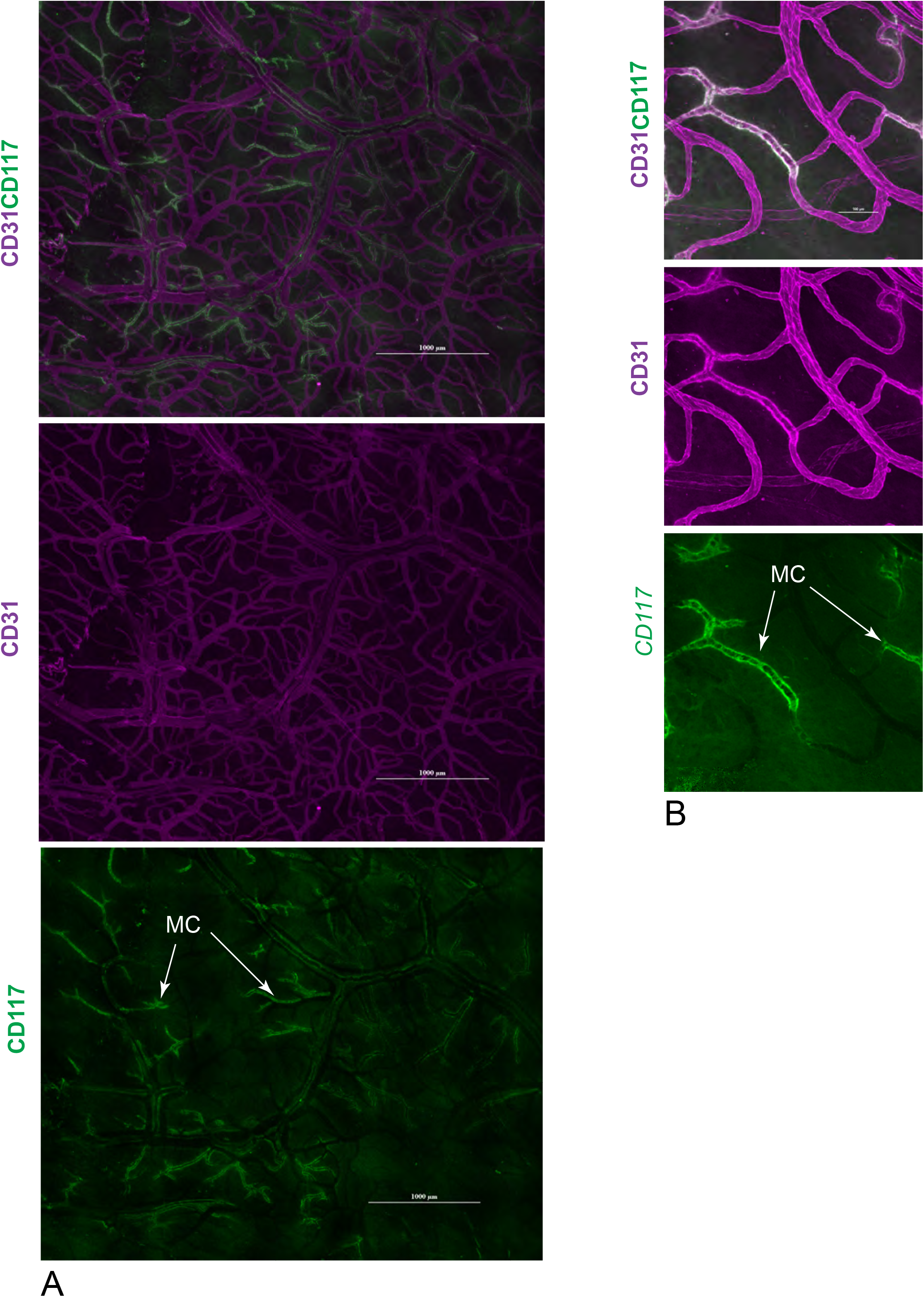
CD117 staining shows close association of CD117+ mast cells with blood vasculature throughout the tissue. (A) Overview of mast cell distribution in monkey dura in lateral area. (B) Close up view of CD117+ mastocytes aligned along a venule wall. Scale bars are 1000 µm on (A) and 100 µm on (B).

### Dura in NPH models of tauopathy and synucleinopathy, with implications for brain clearance

Meninges, and specifically lymphatics in dura, have been implicated in clearance of macromolecules from the brain.^28,29,50^ Studies in rodent models showed that if the meningeal lymphatic system is compromised, this exacerbates neurodegenerative diseases such as Alzheimer’s (AD) and Parkinson’s (PD).^51–54^ However, information about meningeal structure and function in primates is very limited. Therefore, we used our imaging protocols to analyze dura preserved from established NHP models of tauopathy and synucleinopathy during necropsies for other studies. Dura from 2 animals of 12 and 14 years old with induced tauopathy and from age matched controls was probed with antibodies against tau, amyloid (4G8), CD31, and PDPN. Surprisingly, we found significant amyloid deposition at various areas of dura in all 3 animals, despite no amyloidosis inducing treatment. 4G8 immunoreactivity was evident in the form of amyloid deposits throughout the tissue and along the arterial walls as found in cerebral amyloid angiopathy (CAA). Panoramic images showed the strongest staining in the area of falx and around MMA (Figure 14). These amyloid deposits usually appear as dense perivascular clusters, as seen in Figure 15. CAA appeared exclusively around arteries without affecting the veins and it was most prominent around MMA and its branches (Figure 16, A and B) and Figure 17. We also observed specific 4G8 immunoreactivity within the lymphatics flanking MMA and arteries (Figure 16, B and C). As seen in these images, amyloid appears to be within the lymphatic lumen, and not along the vessel wall. Deposition along and around the arterial vessels, and not inside it, is apparent even for small arteries. Unlike the lateral lymphatics around MMA, the lymphatics associated with SSS were largely negative for 4G8 immunoreactivity, which is clearly seen in Figure 15. Staining with anti-tau antibody showed diffuse signal throughout the meningeal tissue and immunoreactivity at the walls of arteries, specifically MMA. We found that tau signal is associated with the arterial wall but not with lymphatics, unlike amyloid (Figure 18). Analysis of dura from the animals treated with α-synuclein fibrils showed the same staining pattern as for amyloid but was less defined (data not shown).

**Figure 14.**
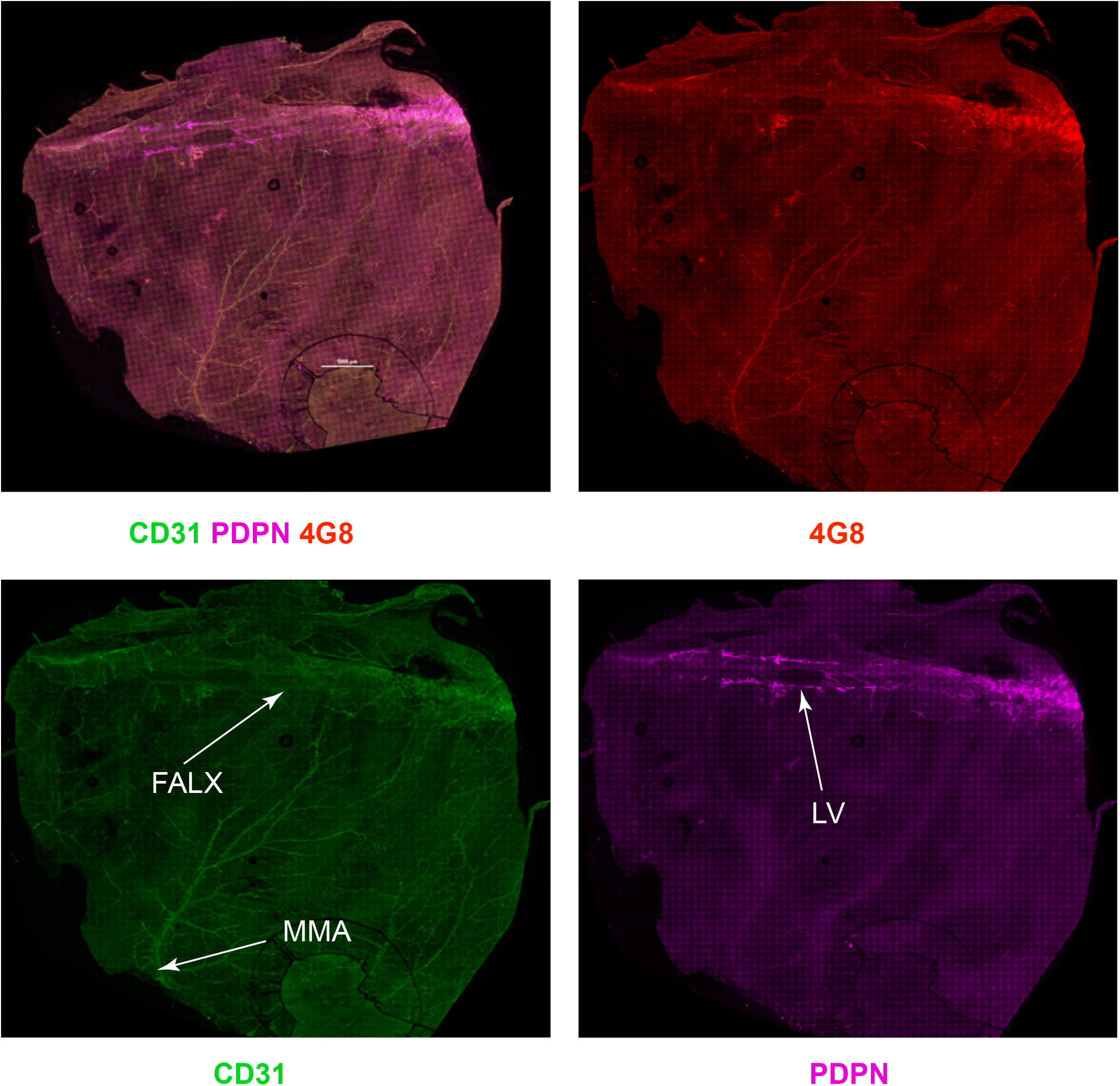
Whole-mount meninges from the tauopathy model. Meninges from the model of tauopathy were probed with CD31, PDPN and 4G8 antibodies. Individual channels and overlay imaging of the lateral dura and the falx area show distribution of the blood vessels with CD31 antibody, lymphatic vessels are seen around flax area as visualized by anti-PDPN antibody. Note that there are PDPN+ vessels flanking MMA and branching arteries not seen at this magnification. 4G8 detects diffuse Aβ staining throughout the tissue, in deposits and along arterial vessels. Scale bar is 5000 µm.

**Figure 15.**
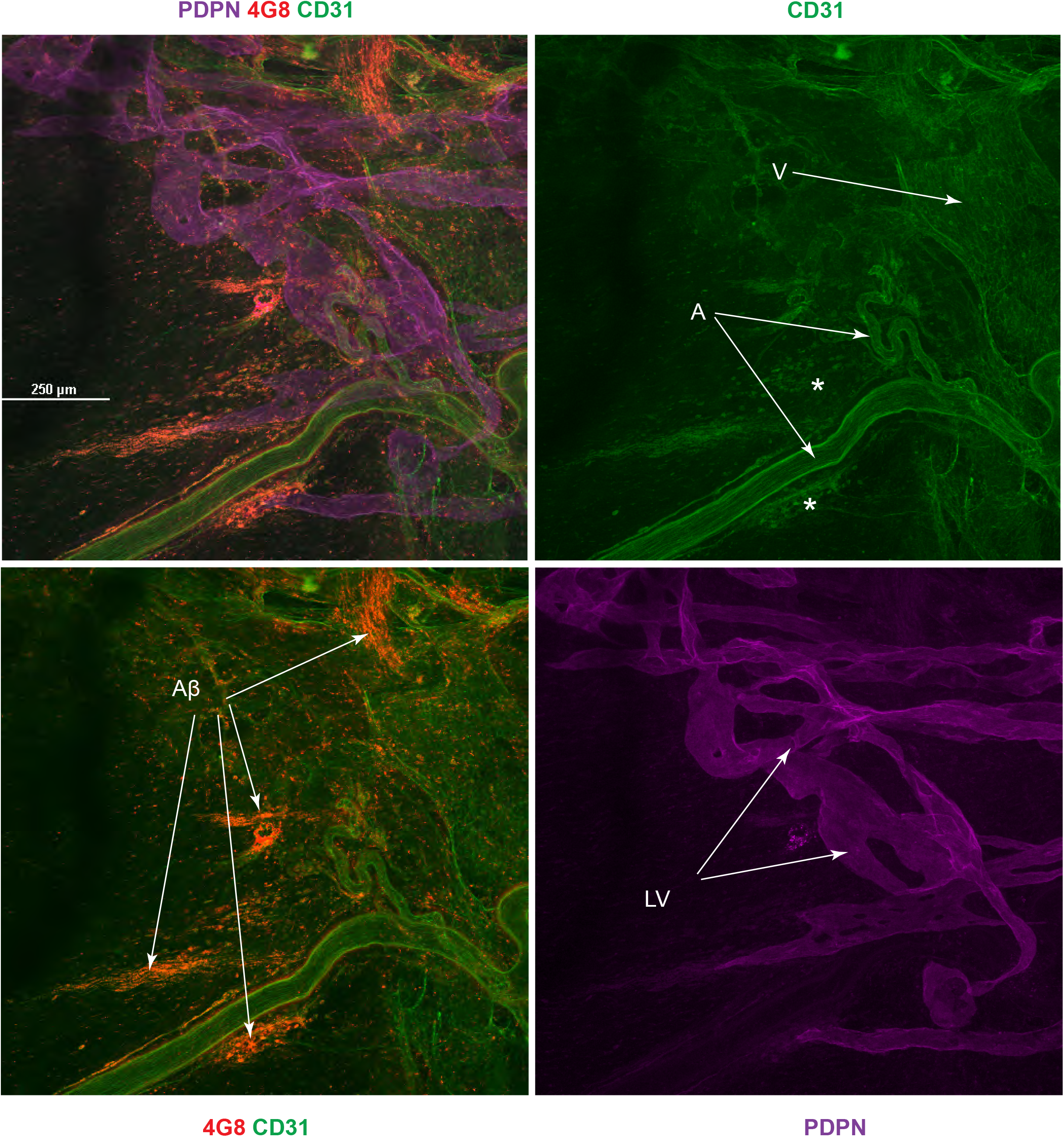
Non-vascular and vascular amyloid deposition in the meninges of the amyloidosis model in the area of SSS. Amyloid deposits without association with blood vessel walls and along arterial walls are clearly seen. There is no amyloid associated with lymphatic vessels around SSS. V – vein, A – artery, LV – lymphatic vessel. Asterisks mark the areas with CD31+ cells. Scale bar is 250 µm.

**Figure 16.**
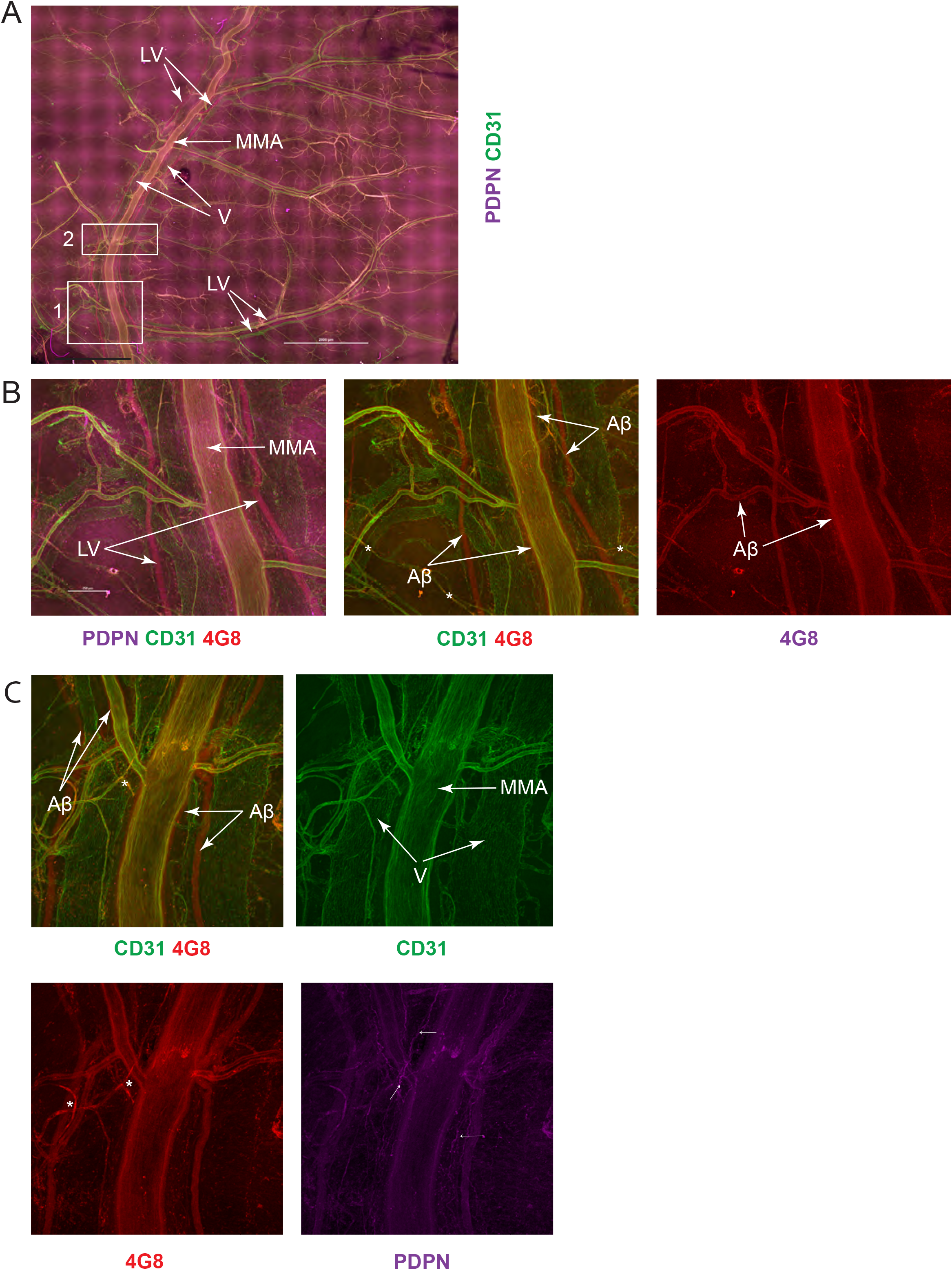
Amyloid is associated with the lumen of lymphatics flanking MMA and seen as CAA around the wall of MMA. (A) MMA is flanked by two lymphatic vessels. (B) and (C) are close up views of the framed areas 1 and 2 on (A) where 4G8 immunoreactivity is clearly seen along MMA and smaller branching arteries and in the lumen of lymphatics. MMA – middle meningeal artery, V – vein, A – artery, LV – lymphatic vessel. Scale bars are 2000 µm on (A).

**Figure 17.**
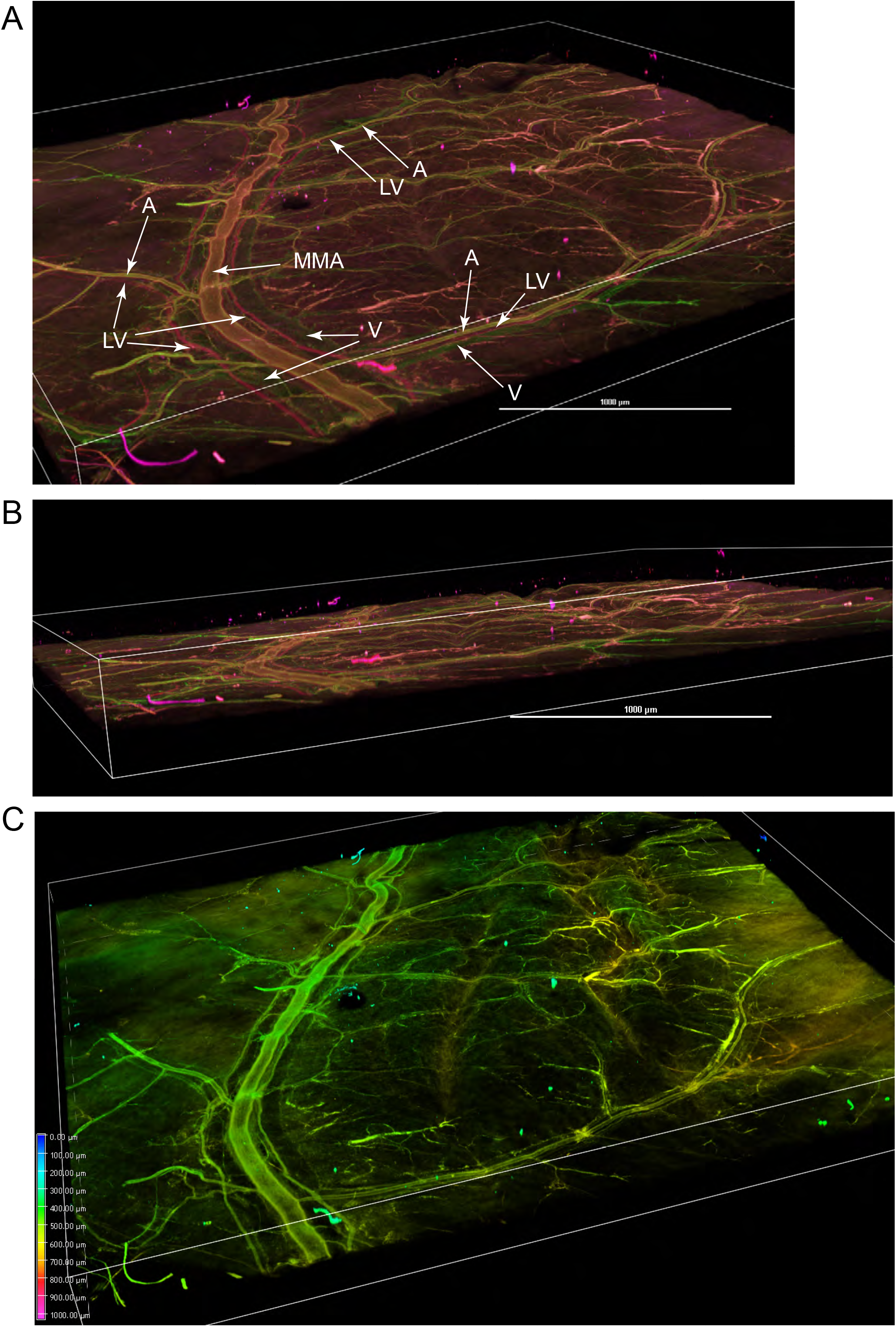
3D rendering of blood and lymphatic vessels in MMA region. Volume projection of the MMA region shown on Figure 16. (A) and (B) show original staining with anti-CD31, anti-PDPN and 4G8 antibodies as seen close to XY and Z view planes. (C) shows the depth heat map demonstrating that the vessels are located on the same plane. MMA – middle meningeal artery, V – vein, A – artery, LV – lymphatic vessel. Scale bars are 1000 µm.

**Figure 18.**
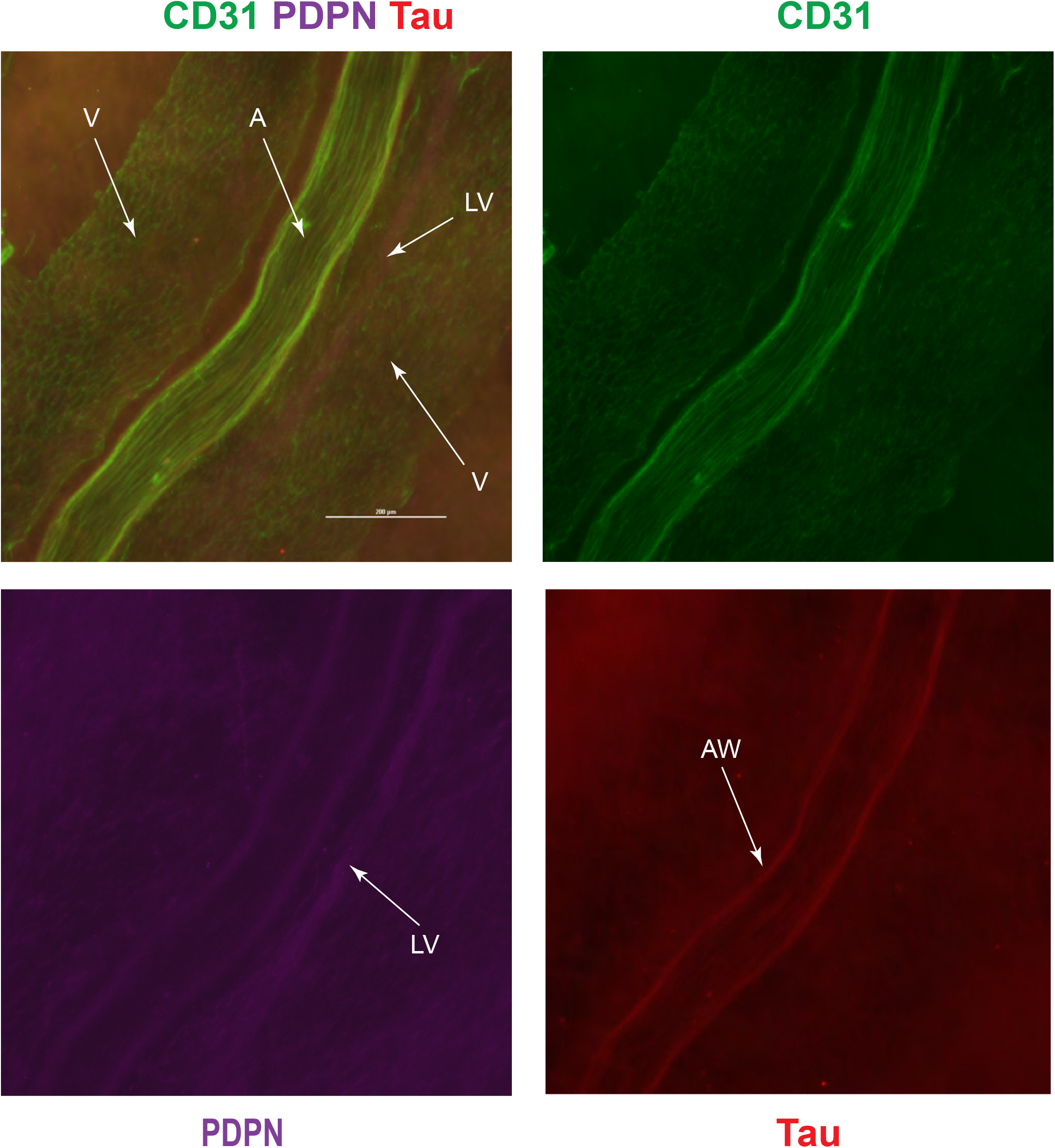
Diffuse Tau immunoreactivity in the dura and around arterial walls. Majority of tau signal was not specific and was diffusely distributed in the dura. Defined signal was observed around arterial walls, especially around MMA. V – vein, A – artery, LV – lymphatic vessel, AW – arterial wall. Scale bars are 200 µm.

## Discussion

We imaged and analyzed 20 whole-mount specimens of dura mater from wild-type Cynomogolus monkeys, 3 specimens of a NHP model of tauopathy (Rhesus), and 4 specimens from a NHP model of synucleinopathy (Cynomogolus). This is the first study to use whole-mount dura preparations of NHP including primate models of neurodegenerative diseases. To date, imaging of whole-mount specimens has been used only for rodent meninges which may not fully replicate human patterns of brain clearance or disease.^28–30,50^

Most information on dural blood supply and anatomy of the major arteries and veins in primates comes from earlier studies using dye injections and gross dissections.^55–57^ Other techniques for live imaging include modalities such as MRI that reveal gross anatomy but lack resolution to identify smaller structures and their spatial relationships. Having a better knowledge of morphological and functional characteristics of blood and lymphatic vasculature in the dura for neurological diseases and surgical applications is clinically important. Meninges are implicated in macromolecular clearance from the brain and brain immunity, which is relevant to many neurological diseases including Alzheimer’s disease.^12,15,17^ In terms of surgical interventions, MMA embolization is an established endovascular approach to management of meningeal tumors and chronic subdural hematomas.^58^ However, there are individual variations in anatomy and anastomoses that pose risk for the procedure and the normal role of MMA is still not well-understood.^59^ MMA enters the skull and branches into anterior and posterior branches. Branching varies among individual animals with middle branch stemming from anterior or posterior branches or absent altogether.^59^ Even less is known about dural venous and lymphatic anatomy, especially in primates.

Our approach for imaging whole-mount cleared tissues provides tools for improved high-resolution imaging of primate dura. Combining clearing and fluorescence based confocal resonance microscopy, we visualized the smallest capillaries and fine detail of vascular anastomoses. We found an extensive vascular meningeal network with dense capillary coverage having cross-sectional diameter of 10-15 μm, larger than blood-brain barrier capillaries. This blood vessel network is much more substantial and denser than observed in rodents. Morphology and staining with αSMA showed that dural blood vessels consist of primarily veins and venules. MMA and branches which supply the dura mater are easily identified. Anatomically, MMA in NHP is very similar to human, including the presence of flanking veins and lymphatic draining channels. Typical vascular anatomy in the dura is that veins follow the arteries. It has been earlier suggested that this proximity may relate to possible communication between arteries and veins without a capillary network.^59^ We consistently observed that accompanying veins are substantially larger than the arteries themselves. This is important to consider, provided that the venous network is very extensive and can be possibly involved in CSF outflow and macromolecular efflux from dura. Non-endothelial CD31+ cells that we observed throughout the tissue are likely hematopoietic cells. CD31, or PECAM-1, is platelet-endothelial cell adhesion protein that was initially identified from these cells and play immunoregulatory roles. While it is still considered as a specific marker for endothelial cells, it is expressed by leukocytes, monocytes, and plasma cells.^39^

One of the questions in the field which attracted high interest but has not been completely resolved is whether lymphatic vessels described in rodent meninges are anatomically and physiologically similar to those in higher mammals and especially in primates. Their presence has been confirmed in monkeys and humans but imaging modalities such as MRI or routine IHC of tissue sections has provided limited knowledge about this vasculature in larger mammals.^16,33,35,36^ Smaller size and minimal thickness proved to be advantageous for studies of meningeal lymphatic system in rodent models because it was possible to prepare whole-mount specimens.

Specimens prepared in this manner allow a panoramic view of tissue with respect to blood and lymphatic vessels.^28,29^ Here we expand upon that work with a methodology for clearing and imaging NHP meninges.

In some cases, discrimination of lymphatics from blood vessels is not straightforward because lymphatics and venous endothelium share some markers.^60^ Examination of collective expression of the panel of the markers LYVE-1, PDPN, CD31, and Prox-1 is usually needed. PDPN is one the specific markers not expressed in blood vessels but expressed in macrophages.^61,62^ Expression of these proteins also varies within the lymphatic system itself depending on the vessel type. Capillaries usually have LYVE-1+/PDPN+ phenotype with LYVE-1 expression decreasing with a transition to the collecting phenotype.^63^ Expression and 5’-Nase activity as well as LA10 antibody have been used for identification of lymphatics.^43,45^ We probed our specimens with a panel of lymphatic markers. Overall, the lymphatic system in the dura of NHP resembles the lymphatic system described in rodents where the vascular structures positive for lymphatic markers were found running along SSS, MMA, and at the skull base.^28–30,50^ We found PDPN+ vessels in the same anatomical locations as in rodents but with differences in appearance and surface marker expression. All PDPN+ vessels were CD31 negative. Our observations agree with those of Goodman *et al* who reported the presence of vessels positive for lymphatic markers along SSS region of human dura and pointed at differences from meningeal lymphatics in mice. ^33^ In their work, the authors performed immunohistochemistry (IHC) on coronal sections of paraffin embedded human dura containing SSS. Similar to our results, the authors observed vessels positive for PDPN and negative for CD31 and LYVE-1 (CD31-/PDPN+/LYVE-1-). They also described another vessel type, which was positive for both LYVE-1 and PDPN+ (CD31-/PDPN+/LYVE-1+). However, we did not identify PDPN+/LYVE-1+ vasculature in our specimens. Occasional faint LYVE signal was observed associated with PDPN+ vessels proximal to SSS, but only with larger pinhole size over 6 AU. These imaging settings most likely do not detect LYVE-1 expression that has physiological significance. PDPN+ vessels flanking MMA and in tentorium were negative for LYVE-1.

We were not able to demonstrate Prox-1 expression in any PDPN+ and LYVE+ structures after testing a variety of anti-Prox-1 antibodies that were confirmed to detect monkey and human antigens.^30,35,36,50^ However, we can’t conclude that Prox-1 is not expressed in those structures because lack of clear signal may reflect an undefined limitation of imaging. Prox-1 is a transcription factor localized in the nucleus and is seen as punctate signal on lymphatic structures. Detection of the discrete nuclear signals usually requires objectives with higher magnification combined with high numerical aperture (NA). In our study, range of acquisition depths was 700-1600 µm, which is only possible with air long working distance lenses. However, long working distance inversely correlates with NA, i.e. resolving power of the lens and intensity of the detected signal.^64^ It is possible that even with the highest NA available to us with long working distance air lenses, the magnification was not sufficient to detect nuclear signal or conversely NA of the highest magnification 40x lens did not provide sufficient resolution.

The PDPN+/LYVE-1-phenotype we report here for monkey dura is different from LYVE-1 positive vessels in the area of SSS and MMA reported previously in murine studies and thought to have capillary phenotype.^28,29^ Goodman *et al* suggested that PDPN+/LYVE-1-phenotype without αSMA expression may be indicative for pre-collector lymphatic vessels. However, in that study the authors used coronal sections, which allow only visualization of the lumen and provide limited morphological information about the vessel appearance and branching. On our whole-mount preparations we were able to assess these morphological features. Most of the vessels around SSS were blind ended as typical for lymphatic capillaries, and therefore, most likely are not pre-collectors. The network of PDPN+ vessels in the SSS area is substantially more extensive than in rodents, with a wide range of vessel diameters, which Goodman *et al* also reported in the range of 19-140 µm. Paired lymphatic vessels flanking the middle meningeal artery and veins have drastically different appearance, always with two vessels on each side of the artery, branching only at the points of anastomoses into smaller vessels that continue to follow the smaller arterial vessels. In another study by Absinta *et al* lymphatic vessel diameters were reported in a wide range of 7-842 µM in marmoset sections.^35^ Similarly to our data, they found the highest density of lymphatic vessels in the area of SSS. The lumen diameter in our specimens varied from 20 to 300 µM in certain sites. In mice, meningeal lymphatic diameter was typically reported as 20-30 µm. It is possible that some vessels collapse during tissue dissection and processing and this adds to size variability. While veins share some endothelial markers with lymphatics, our imaging showed a clear distinction between veins and lymphatics. First, lymphatic PDPN+ vessels were consistently negative for CD31 endothelial marker that visualizes blood vasculature. Second, we also observed red blood cells (RBC) even in smaller arterioles but never in PDPN+ vessels. Therefore, these vessels are not blood vasculature. Possibly, these vessels should not be defined as lymphatics and they are special vascular structures unique to CNS environment.

In terms of relative roles of clearance in health and disease, the contribution of perivascular efflux along arteries versus intraluminal clearance in veins and lymphatics remains to be fully worked out. Our data suggest that multiple mechanisms of clearance are involved, including both sagittal and lateral lymphatic structures in the meninges. This implies that transport of solutes through the entire arachnoid membrane surrounding the brain is likely important as a gatekeeper to widespread transport.

Deposition of amyloid in the dura of patients with Alzheimer’s disease occurs both in the form of diffuse deposits and vascular amyloid pathology. Moreover, CAA of meningeal vessels in Alzheimer’s disease occurs earlier than in intracerebral vessels.^8,65,66^ Similar pathology in meninges is demonstrated in 5xFAD mice, and aggravated by ablation of meningeal lymphatics suggesting a contribution to amyloid clerance.^53^ In our work, amyloid was probed with 4G8, which displays broad reactivity to amyloid species. In NHP dura, we observed distinctive CAA around arteries and amyloid deposits in both control and tauopathy models in aged NHP as early as 14 years of age. This finding points at very early events that cause amyloid deposition specifically in the dura, which possibly implicates inefficiency in Aβ clearance. In prior studies of aging in rhesus monkeys, CAA of the cerebral vessels was not detectable in any animal under the age of 20 years.^67,68^ Similar to rhesus monkeys, cynomolgus monkeys manifest CAA after 20 years of age, and there is a positive correlation between parenchymal plaques and CAA.^67– 71^ As with findings reported by Goodman *et al*, we did not find Aβ deposition along lymphatic vessel walls or within the lumen in the area of SSS. The SSS lymphatics were completely devoid from 4G8 immunoreactivity. In that study the MMA lymphatics were not analyzed. We found that staining with 4G8 antibody produces strong positive signal in lymphatics flanking MMA which is consistent with Aβ inside the lumen. This is especially evident when compared to staining of smaller arteries where Aβ is seen deposited along the vessel walls leaving the lumen clear. We also found meningeal perivascular localization of α-synuclein in a NHP model of synucleinopathy similar to the localization of Aβ. Localization of 4G8 positive areas was well-defined compared to more diffuse meningeal signal for tau, which was observed throughout the dura. More defined tau immunoreactivity was observed along arterial walls, especially MMA, but without signal in the flanking lymphatics. This also contrasts with Aβ presence in those vessels and suggests that dural clearance pathways in primates for Aβ and tau are distinct but complementary.

## Methods

### Animals and tissue preparation

All experiments were approved by the Institutional Animal Care and Use Committees at the sites of animal procedures and necropsies: Rush University Medical Center, University of Illinois at Chicago and University of California, Davis. For tissue collection animals were sacrificed by transcardiac perfusion with PBS or 4% PFA solution under anesthesia. If perfused with PBS, the tissues were fixed overnight in 4% PFA. Dura was dissected and processed for staining. For experiments related to tauopathy and synucleinopathy, we used tissues from the monkey models generated as previously described for other studies.^37,38^

### Immunofluorescence

Tissues were washed in PBS, incubated overnight with blocking solution containing 1% bovine serum albumin (BSA) and 5% of the appropriate serum and then with the primary antibody at 4 °C overnight. After incubation with primary antibodies, tissues were washed incubated with secondary antibodies. After washing in PBS, tissues were clarified using modified BABB clearing protocol and mounted with Cytoseal 60 or 260 (Richard-Allan Scientific) and dried overnight.

### Imaging

Images were acquired on Nikon A1 confocal microscope with different objectives and resonant or Galvano scanners, depending on the experiment. Imaging of larger fields was performed using high-speed resonant scanner and long working distance 10x and 20x lenses that allowed reasonable acquisition times. Imaging of the regions of interest with higher magnification was done using Galvano scanner with 10x, 20x and 40x objectives. (name N/A air). Elements (Nikon) or Imaris software was used for processing and analysis of images. Imaging algorithms were tile scans with z-stack acquisition where z-section thickness varied from 0.5 to 8 µm depending on the experiment. When needed, maximum intensity or 3D projections were generated.

## References

1. Weller, R. O., Sharp, M. M., Christodoulides, M., Carare, R. O. & Møllgård, K. The meninges as barriers and facilitators for the movement of fluid, cells and pathogens related to the rodent and human CNS. Acta Neuropathol. (Berl.) 1–23 (2018) doi:10.1007/s00401-018-1809-z.

2. Rua, R. & McGavern, D. B. Advances in Meningeal Immunity. Trends Mol. Med. 24, 542–559 (2018).

3. Derecki, N. C. et al. Regulation of learning and memory by meningeal immunity: a key role for IL-4. J. Exp. Med. 207, 1067–1080 (2010).

4. Coles, J. A., Myburgh, E., Brewer, J. M. & McMenamin, P. G. Where are we? The anatomy of the murine cortical meninges revisited for intravital imaging, immunology, and clearance of waste from the brain. Prog. Neurobiol. 156, 107–148 (2017).

5. Benakis, C., Llovera, G. & Liesz, A. The meningeal and choroidal infiltration routes for leukocytes in stroke. Ther. Adv. Neurol. Disord. 11, 1756286418783708 (2018).

6. Arac, A., Grimbaldeston, M. A., Galli, S. J., Bliss, T. M. & Steinberg, G. K. Meningeal Mast Cells as Key Effectors of Stroke Pathology. Front. Cell. Neurosci. 13, (2019).

7. Alves de Lima, K., Rustenhoven, J. & Kipnis, J. Meningeal Immunity and Its Function in Maintenance of the Central Nervous System in Health and Disease. Annu. Rev. Immunol. 38, 597–620 (2020).

8. Takeda, S., Yamazaki, K., Miyakawa, T. & Onda, K. Cerebral amyloid angiopathy initially occurs in the meningeal vessels. Neuropathol. Off. J. Jpn. Soc. Neuropathol. 37, 502–508 (2017).

9. Levy, D., Labastida-Ramirez, A. & MaassenVanDenBrink, A. Current understanding of meningeal and cerebral vascular function underlying migraine headache. Cephalalgia Int. J. Headache 39, 1606–1622 (2019).

10. Mesquita, S. D., Fu, Z. & Kipnis, J. The Meningeal Lymphatic System: A New Player in Neurophysiology. Neuron 100, 375–388 (2018).

11. Derk, J., Jones, H. E., Como, C., Pawlikowski, B. & Siegenthaler, J. A. Living on the Edge of the CNS: Meninges Cell Diversity in Health and Disease. Front. Cell. Neurosci. 15, 245 (2021).

12. Rua, R. et al. Infection drives meningeal engraftment by inflammatory monocytes that impairs CNS immunity. Nat. Immunol. 20, 407–419 (2019).

13. Yanev, P. et al. Impaired meningeal lymphatic vessel development worsens stroke outcome: J. Cereb. Blood Flow Metab. (2019) doi:10.1177/0271678X18822921.

14. Wojciechowski, S. et al. Developmental Dysfunction of the Central Nervous System Lymphatics Modulates the Adaptive Neuro-Immune Response in the Perilesional Cortex in a Mouse Model of Traumatic Brain Injury. Front. Immunol. 11, 559810 (2021).

15. Bolte, A. C. et al. Meningeal lymphatic dysfunction exacerbates traumatic brain injury pathogenesis. bioRxiv 817023 (2019) doi:10.1101/817023.

16. Visanji, N. P., Lang, A. E. & Munoz, D. G. Lymphatic vasculature in human dural superior sagittal sinus: Implications for neurodegenerative proteinopathies. Neurosci. Lett. 665, 18–21 (2018).

17. Louveau, A., Mesquita, S. D. & Kipnis, J. Lymphatics in Neurological Disorders: A neuro-lympho-vascular Component of Multiple Sclerosis and Alzheimer’s Disease. Neuron 91, 957–973 (2016).

18. Protasoni, M. et al. The collagenic architecture of human dura mater. J. Neurosurg. 114, 1723–1730 (2011).

19. Mecheri, B., Paris, F. & Lübbert, H. Histological investigations on the dura mater vascular system of mice. Acta Histochem. 120, 846–857 (2018).

20. Ma, T., Wang, F., Xu, S. & Huang, J. H. Meningeal immunity: Structure, function and a potential therapeutic target of neurodegenerative diseases. Brain. Behav. Immun. 93, 264–276 (2021).

21. Shukla, V., Hayman, L. A. & Taber, K. H. Adult cranial dura II: venous sinuses and their extrameningeal contributions. J. Comput. Assist. Tomogr. 27, 98–102 (2003).

22. Nabeshima, S., Reese, T. S., Landis, D. M. & Brightman, M. W. Junctions in the meninges and marginal glia. J. Comp. Neurol. 164, 127–169 (1975).

23. Saunders, N. R., Habgood, M. D., Møllgård, K. & Dziegielewska, K. M. The biological significance of brain barrier mechanisms: help or hindrance in drug delivery to the central nervous system? F1000Research 5, (2016).

24. Saunders, R. L. & Bell, M. A. X-ray microscopy and histochemistry of the human cerebral blood vessels. J. Neurosurg. 35, 128–140 (1971).

25. Ichimura, T., Fraser, P. A. & Cserr, H. F. Distribution of extracellular tracers in perivascular spaces of the rat brain. Brain Res. 545, 103–113 (1991).

26. Rai, R., Iwanaga, J., Shokouhi, G., Oskouian, R. J. & Tubbs, R. S. The Tentorium Cerebelli: A Comprehensive Review Including Its Anatomy, Embryology, and Surgical Techniques. Cureus (2018) doi:10.7759/cureus.3079.

27. Weller, R. O., Sharp, M. M., Christodoulides, M., Carare, R. O. & Møllgård, K. The meninges as barriers and facilitators for the movement of fluid, cells and pathogens related to the rodent and human CNS. Acta Neuropathol. (Berl.) 1–23 (2018) doi:10.1007/s00401-018-1809-z.

28. Louveau, A. et al. Structural and functional features of central nervous system lymphatics. Nature 523, 337–341 (2015).

29. Aspelund, A. et al. A dural lymphatic vascular system that drains brain interstitial fluid and macromolecules. J. Exp. Med. 212, 991–999 (2015).

30. Jung, E. et al. Development and Characterization of A Novel Prox1-EGFP Lymphatic and Schlemm’s Canal Reporter Rat. Sci. Rep. 7, (2017).

31. Miura, M., Kato, S. & von Lüdinghausen, M. Lymphatic drainage of the cerebrospinal fluid from monkey spinal meninges with special reference to the distribution of the epidural lymphatics. Arch. Histol. Cytol. 61, 277–286 (1998).

32. Welch, K. & Pollay, M. Perfusion of particles through arachnoid villi of the monkey. Am. J. Physiol. 201, 651–654 (1961).

33. Goodman, J. R., Adham, Z. O., Woltjer, R. L., Lund, A. W. & Iliff, J. J. Characterization of dural sinus-associated lymphatic vasculature in human Alzheimer’s dementia subjects. Brain. Behav. Immun. 73, 34– 40 (2018).

34. Albayram, M. S. et al. Non-invasive MR imaging of human brain lymphatic networks with connections to cervical lymph nodes. Nat. Commun. 13, 203 (2022).

35. Absinta, M. et al. Human and nonhuman primate meninges harbor lymphatic vessels that can be visualized noninvasively by MRI. eLife 6, e29738 (2017).

36. Antila, S. et al. Development and plasticity of meningeal lymphatic vessels. J. Exp. Med. jem.20170391 (2017) doi:10.1084/jem.20170391.

37. Chu, Y. et al. Intrastriatal alpha-synuclein fibrils in monkeys: spreading, imaging and neuropathological changes. Brain 142, 3565–3579 (2019).

38. Beckman, D. et al. A novel tau-based rhesus monkey model of Alzheimer’s pathogenesis. Alzheimers Dement. 17, 933–945 (2021).

39. Woodfin, A., Voisin, M.-B. & Nourshargh, S. PECAM-1: A Multi-Functional Molecule in Inflammation and Vascular Biology. Arterioscler. Thromb. Vasc. Biol. 27, 2514–2523 (2007).

40. Aguilar-Pineda, J. A. et al. Vascular smooth muscle cell dysfunction contribute to neuroinflammation and Tau hyperphosphorylation in Alzheimer disease. iScience 24, 102993 (2021).

41. Brezovakova, V. & Jadhav, S. Identification of Lyve-1 positive macrophages as resident cells in meninges of rats. J. Comp. Neurol. 528, 2021–2032 (2020).

42. Podgrabinska, S. et al. Molecular characterization of lymphatic endothelial cells. Proc. Natl. Acad. Sci. 99, 16069–16074 (2002).

43. Ji, R.-C., Qu, P. & Kato, S. Application of a New 5′-Nase Monoclonal Antibody Specific for Lymphatic Endothelial Cells. Lab. Invest. 83, 1681–1683 (2003).

44. Ji, R.-C. & Kato, S. Lymphatic Network and Lymphangiogenesis in the Gastric Wall. J. Histochem. Cytochem. 51, 331–338 (2003).

45. Ezaki, T. et al. Production of two novel monoclonal antibodies that distinguish mouse lymphatic and blood vascular endothelial cells. Anat. Embryol. (Berl.) 211, 379–393 (2006).

46. Cueni, L. N. & Detmar, M. The Lymphatic System in Health and Disease. Lymphat. Res. Biol. 6, 109–122 (2008).

47. Ulvmar, M. H. & Mäkinen, T. Heterogeneity in the lymphatic vascular system and its origin. Cardiovasc. Res. 111, 310–321 (2016).

48. Sato, T., Konishi, H., Tamada, H., Nishiwaki, K. & Kiyama, H. Morphology, localization, and postnatal development of dural macrophages. Cell Tissue Res. (2021) doi:10.1007/s00441-020-03346-y.

49. Vukman, K. V., Försönits, A., Oszvald, Á., Tóth, E. Á. & Buzás, E. I. Mast cell secretome: Soluble and vesicular components. Semin. Cell Dev. Biol. 67, 65–73 (2017).

50. Ahn, J. H. et al. Meningeal lymphatic vessels at the skull base drain cerebrospinal fluid. Nature 572, 62–66 (2019).

51. Wen, Y.-R., Yang, J.-H., Wang, X. & Yao, Z.-B. Induced dural lymphangiogenesis facilities soluble amyloid-beta clearance from brain in a transgenic mouse model of Alzheimer’s disease. Neural Regen. Res. 13, 709–716 (2018).

52. Wang, L. et al. Deep cervical lymph node ligation aggravates AD-like pathology of APP/PS1 mice. Brain Pathol. 0,.

53. Da Mesquita, S. et al. Functional aspects of meningeal lymphatics in ageing and Alzheimer’s disease. Nature (2018) doi:10.1038/s41586-018-0368-8.

54. Zou, W. et al. Blocking meningeal lymphatic drainage aggravates Parkinson’s disease-like pathology in mice overexpressing mutated α-synuclein. Transl. Neurodegener. 8, 7 (2019).

55. Shabo, A. L., Abbott, M. M. & Maxwell, D. S. The response of the arachnoid villus to an intracisternal injection of autogenous brain tissue. An electron microscopic study in the macaque monkey. Neurology 19, 724–734 (1969).

56. Rutka, J. T. et al. An Ultrastructural and Immunocytochemical Analysis of Leptomeningeal and Meningioma Cultures. J. Neuropathol. Exp. Neurol. 45, 285–303 (1986).

57. Kida, S., Yamashima, T., Kubota, T., Ito, H. & Yamamoto, S. A light and electron microscopic and immunohistochemical study of human arachnoid villi. J. Neurosurg. 69, 429–435 (1988).

58. Martínez, J. L. et al. The Middle Meningeal Artery: Branches, Dangerous Anastomoses, and Implications in Neurosurgery and Neuroendovascular Surgery. Oper. Neurosurg. 22, 1–13 (2022).

59. Falk, D. & Nicholls, P. Meningeal arteries in rhesus macaques (Macaca mulatta): Implications for vascular evolution in anthropoids. Am. J. Phys. Anthropol. 89, 299–308 (1992).

60. Kong, L.-L. et al. The optimum marker for the detection of lymphatic vessels (Review). Mol. Clin. Oncol. 7, 515–520 (2017).

61. Bieniasz-Krzywiec, P. et al. Podoplanin-Expressing Macrophages Promote Lymphangiogenesis and Lymphoinvasion in Breast Cancer. Cell Metab. 30, 917-936.e10 (2019).

62. JCI Insight - Monocyte upregulation of podoplanin during early sepsis induces complement inhibitor release to protect liver function. https://insight.jci.org/articles/view/134749.

63. Weber, E. et al. Lymphatic Collecting Vessels in Health and Disease: A Review of Histopathological Modifications in Lymphedema. Lymphat. Res. Biol. (2022) doi:10.1089/lrb.2021.0090.

64. North, A. J. Seeing is believing? A beginners’ guide to practical pitfalls in image acquisition. J. Cell Biol. 172, 9–18 (2006).

65. Chen, X. et al. Cerebral amyloid angiopathy is associated with glymphatic transport reduction and time-delayed solute drainage along the neck arteries. Nat. Aging 2, 214–223 (2022).

66. Kovacs, G. G. et al. Dura mater is a potential source of Aβ seeds. Acta Neuropathol. (Berl.) 131, 911–923 (2016).

67. Uno, H. et al. Cerebral amyloid angiopathy and plaques, and visceral amyloidosis in aged macaques. Neurobiol. Aging 17, 275–281 (1996).

68. Emborg, M. E. Nonhuman Primate Models of Neurodegenerative Disorders. ILAR J. 58, 190–201 (2017).

69. Martin, L. J. et al. Amyloid precursor protein in aged nonhuman primates. Proc. Natl. Acad. Sci. 88, 1461– 1465 (1991).

70. Teil, M., Arotcarena, M.-L. & Dehay, B. A New Rise of Non-Human Primate Models of Synucleinopathies. Biomedicines 9, 272 (2021).

71. Uno, H. & Walker, L. C. The age of biosenescence and the incidence of cerebral beta-amyloidosis in aged captive rhesus monkeys. Ann. N. Y. Acad. Sci. 695, 232–235 (1993).

